# Computational and causal dissociation in primate striatum during value-guided decision-making

**DOI:** 10.64898/2026.07.29.741388

**Authors:** Ji-Woo Lee, Min-Seo Kim, Seong-Hwan Hwang, Hyoung F. Kim

## Abstract

How neural representations carrying similar information across brain regions are translated into behavioral control remains a fundamental question in neuroscience. In the primate striatum, whether subregions make overlapping or dissociable computational and causal contributions to value-guided decisions is unresolved. Here, we show that macaque striatal subregions make dissociable, modality-specific contributions: the putamen is required for tactile-value decisions, the caudate for visual-value decisions, and the ventral striatum for both. Despite broadly shared value encoding, causal necessity is predicted not by the strength of single-neuron value discrimination or population-level decoding, but by neural geometry–behavior coupling. These findings reveal a functional organization of the primate striatum and identify neural geometry–behavior coupling as a key determinant of behavioral control.

## Main text

Value-guided decisions require the brain to transform sensory information into action. The striatum is widely implicated in this transformation, yet whether its subregions—the caudate, putamen, and ventral striatum—serve functionally distinct or overlapping roles remains unresolved ^1–3^.

Two contrasting views dominate the field. One emphasizes functional specialization, including the actor–critic framework ^4–6^ and the dissociation between goal-directed and habitual behavior, mainly from human fMRI and rodent studies ^7–11^. In contrast, recent non-human primate studies suggest functional overlap, citing shared inputs and similar neural responses across subregions ^10,12–14^. This raises a key question: do striatal subregions perform shared computations or make distinct contributions to behavior?

Several limitations constrain current understanding. Most studies rely on single sensory modalities, particularly vision in primate studies, potentially obscuring modality-specific computations across striatal subregions during decision-making. Additionally, single-neuron analyses may average heterogeneous activity and obscure the distinct structure of neural computation ^15,16^. Population-level approaches can overcome this limitation by revealing latent neural geometry beyond single-neuron analyses ^17–22^. However, even when population-level analyses reveal distinct neural representations across regions, they do not indicate which representations are causally recruited for behavioral control. Linking neural representations to behavior requires direct perturbation, without which the functional roles of striatal subregions remain unresolved.

Here, we demonstrate that striatal subregions make dissociable computational and causal contributions to visual and tactile value-guided decisions. Crucially, causal necessity is determined not by value discrimination strength, but by neural geometry–behavior coupling underlying value generalization. Our results identify neural geometry–behavior coupling as a mechanism linking striatal computations to behavioral control.

### Macaques discriminate and learn tactile and visual values

To test whether monkeys could learn tactile and visual stimulus values, they were trained on two tasks: a tactile value reversal task (T-VRT) using braille patterns and a visual value reversal task (V-VRT) using fractal images (Figs. 1A-B, S1A-D). In T-VRT (Fig. 1A), one of two braille patterns was associated with a reward (Good) whereas the other was not (Bad) within each block. These associations were reversed across blocks to isolate neural responses to tactile value rather than to the stimuli themselves (Fig. 1C). Each T-VRT trial began with the onset of a cue, after which the monkey inserted its left index finger into a hole to explore the braille stimulus. After withdrawal of the finger, a blank delay period of 500-1000 ms followed. The monkey then reinserted its finger to encounter the same stimulus presented during the first insertion. Reaction time (RT) was measured during the second insertion, reflecting the reward value associated with the stimulus encountered during the first insertion. V-VRT had the same structure but used visual instead of tactile stimuli (Fig. 1B).

**Figure 1.**
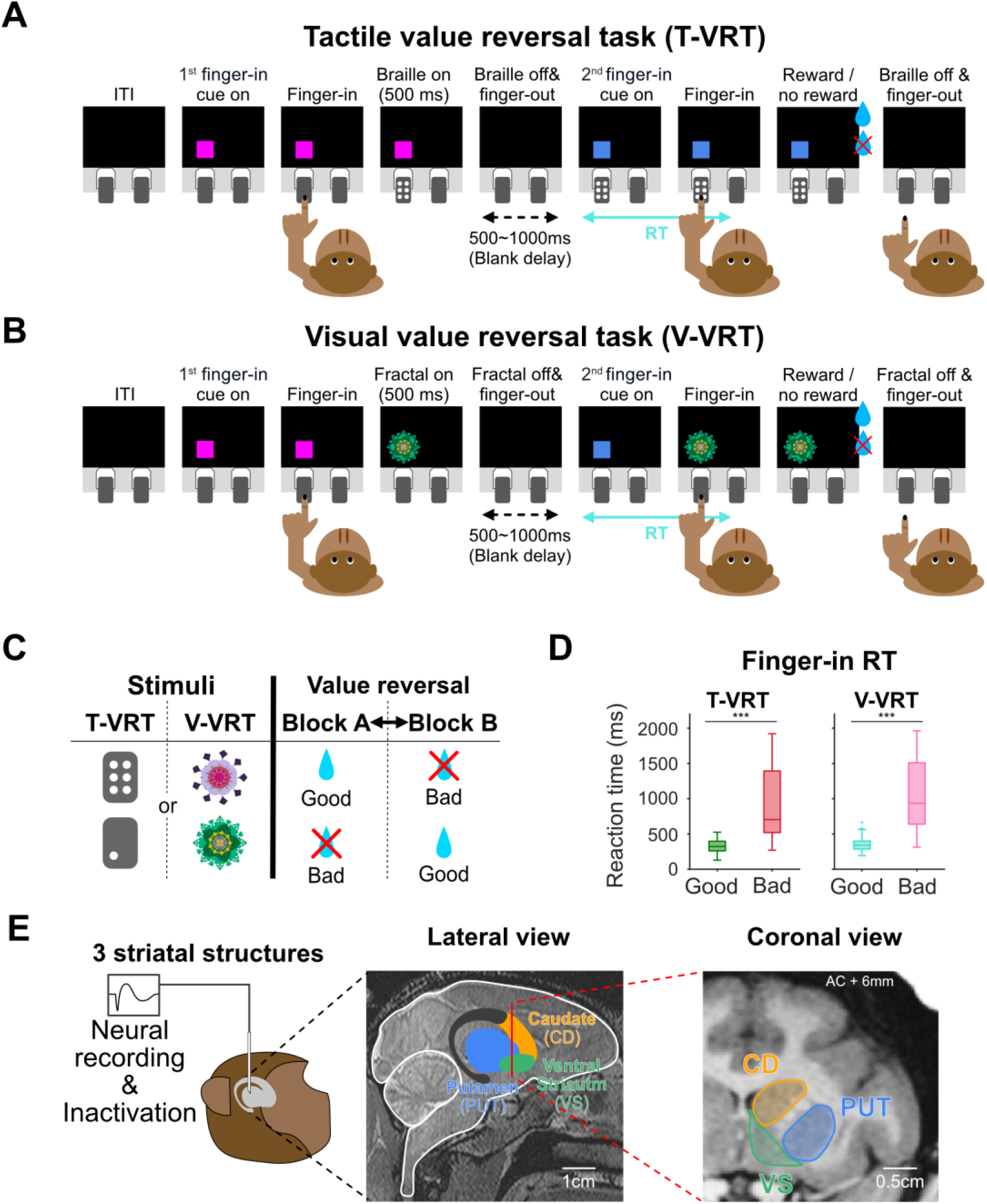
Tactile and visual value learning and electrophysiology across striatal subregions. **(A)** Tactile value reversal task (T-VRT). Schematic of the trial structure. The monkey experiences the same tactile stimulus twice; reaction time (RT) was measured at the second finger insertion. **(B)** Visual value reversal task (V-VRT). The procedure was identical to that in (A), except that fractal images were used as reward-associated visual stimuli. **(C)** Stimulus sets and value reversal. Braille patterns and fractal images used in T-VRT and V-VRT, respectively. Value assignments were reversed across blocks. **(D)** Behavioral performance. Reaction times to good and bad stimuli (776 sessions from two monkeys). Statistical details are provided in Table S2. **(E)** The primate striatum includes the putamen (PUT), caudate (CD), and ventral striatum (VS). Schematic illustration and representative MR images of the primate striatum, showing the putamen, caudate, and ventral striatum in both lateral (*left*) and coronal (*right*) views. Box plot elements: center line, median; box limits, 25th and 75th percentiles; whiskers, 1.5 x IQR; colored crosses, outliers. ***p < 0.0005; n.s., not significant.

In both tasks, monkeys learned the value associations, showing significantly faster RTs for good stimuli than for bad stimuli (T-VRT: 327.07 ± 2.93 vs. 912.04 ± 17.23 ms; V-VRT: 345.34 ± 2.68 vs. 1053.55 ± 16.97 ms) (Figs. 1D, S1E). This was further confirmed in choice trials (Fig. S1F-G), where monkeys reliably discriminated between good and bad stimuli (95.12 ± 0.29% in T-VRT and 98.18 ± 0.15% in V-VRT). This consistent value discrimination across RT and choice behavior enabled us to examine how value information is processed across the putamen, caudate, and ventral striatum at the neuronal level (Fig. 1E).

### Modality-selective and bimodal value neurons across striatal subregions

We previously identified tactile-selective, visual-selective, and bimodal value neurons in the putamen, each encoding value associated with tactile stimuli, visual stimuli, or both, respectively ^23^. Here, we asked whether these neurons extend beyond the putamen and how they are distributed across striatal subregions using single-unit recording (Fig. 1E).

Across striatal subregions, we identified three types of value neurons, with representative examples from the putamen, caudate, and ventral striatum shown in Supplementary figure 2. An example neuron recorded in the putamen responded more strongly to good than to bad tactile stimuli during stimulus presentation but did not discriminate value in V-VRT (Fig. S2A). An example neuron from the caudate showed visual-selective value encoding, responding more strongly to visual stimuli associated with reward during stimulus presentation in V-VRT, but not in T-VRT (Fig. S2B). An example neuron in the ventral striatum encoded both tactile and visual value, exhibiting value discrimination responses during the blank delay period (Fig. S2C).

To systematically characterize these responses, we defined value neurons as those significantly discriminating between good and bad values during stimulus or delay periods in at least one task. These neurons were then classified by modality selectivity, with value encoding assessed across both task periods as previously described (Table S1) ^23^. These results demonstrate the presence of diverse types of value neurons and establish a framework for comparing value representations across striatal subregions at the population level.

### Distribution of modality-selective and bimodal value neurons across striatal subregions

We found 247 (83%), 213 (80%), and 168 (80%) value neurons out of task-relevant neurons in the putamen, caudate, and ventral striatum, respectively. Across all regions, approximately half of the value neurons were bimodal, with the remainder consisting of modality-selective value neurons (Figs. 2A-C and S3A-B). Although the overall proportions were largely comparable across subregions, the caudate contained a significantly different proportions of visual-selective value neurons than the ventral striatum, whereas no significant regional differences were observed for bimodal or tactile-selective value neurons.

**Figure 2.**
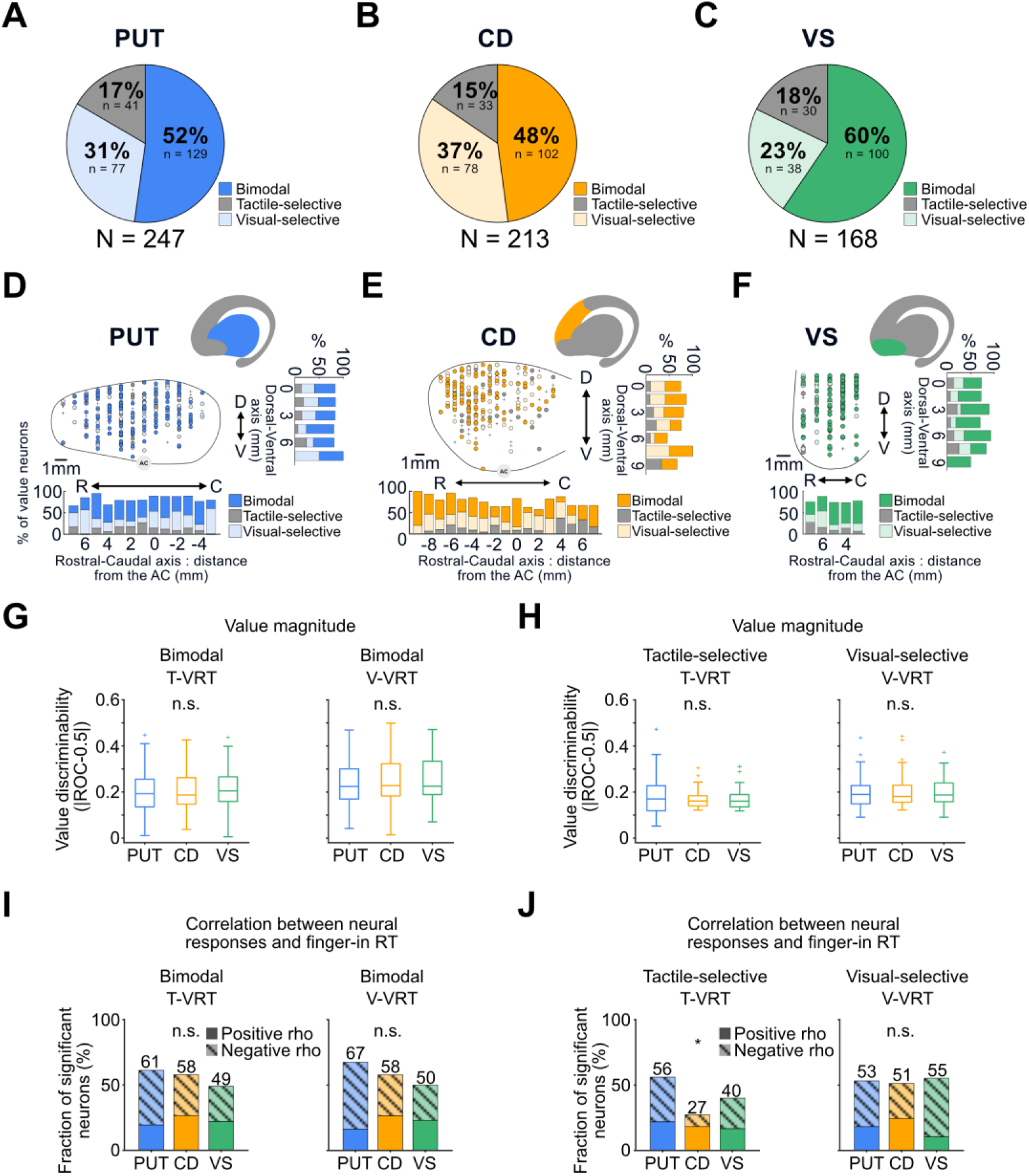
Proportions, anatomical distribution, and single-neuron properties of value neurons across striatal subregions. **(A-C)** Proportions of value neuron types in the putamen **(A)**, caudate **(B)**, and ventral striatum **(C)**. Statistical details are provided in Table S2. **(D-F)** Locations of value neurons. Lateral views show the distribution of value neurons in the putamen **(D)**, caudate **(E)**, and ventral striatum **(F)**. Abbreviations: D, dorsal; V, ventral; R, rostral; C, caudal; AC, anterior commissure. **(G-H)** Value discrimination magnitude measured by ROC analysis. **(G)** Bimodal value neurons. *Left*: T-VRT; *right*: V-VRT. **(H)** Modality-selective value neurons. *Left*: tactile-selective; *right*: visual-selective. **(I-J)** Correlation with reaction time. Proportions of value neurons showing significant correlations between neural responses and subsequent finger-in reaction times (RTs). Bars represent the putamen (blue), caudate (yellow), and ventral striatum (green). Solid bars indicate positive correlations, whereas hatched bars indicate negative correlations. Numbers on each bar denote the total percentage of significant neurons. **(I)** Bimodal value neurons. *Left*: T-VRT; *right*: V-VRT. **(J)** Modality-selective value neurons. *Left*: tactile-selective; *right*: visual-selective. Box plot elements: center line, median; box limits, 25th and 75th percentiles; whiskers, 1.5 x IQR; red crosses, outliers. *p < 0.05; n.s., not significant.

To assess their anatomical organization, we mapped the locations of value neurons across the three regions (Figs. 2D-F and S3C). Value neurons were broadly distributed within each region, with no evidence of clustering or patch-like organization, indicating similar anatomical organization across subregions. We therefore asked whether the properties of value signals differ across striatal subregions at the single-neuron level.

### Comparable value encoding across striatal subregions at the single-neuron level

Averaged responses were similar across striatal subregions for both stimulus- and delay-period value neurons (Figs. S4 and S5A-B). Responses were quantified using peristimulus time histograms (PSTHs), with value discrimination defined as the difference between preferred and non-preferred values, where the preferred value corresponded to the condition (good or bad) eliciting the stronger response in each neuron (Fig. S5A-B).

To further characterize value responses at the individual-neuron level, we analyzed discriminability, duration, and latency. Value discriminability, quantified using receiver operating characteristic (ROC) analysis, did not differ across striatal subregions (Fig. 2G-H). The duration of value encoding was also largely similar across subregions, except that bimodal value neurons in the caudate exhibited a longer duration than those in the putamen in V-VRT (Fig. S5C-D). We next examined value onset latencies. Bimodal and visual-selective value neurons in the caudate, as well as bimodal value neurons in the ventral striatum, responded more rapidly than those in the putamen, whereas tactile-selective value neurons showed no differences (Fig. S5E-F). These results show that single-neuron value responses were largely comparable across striatal subregions, with only a modest bias in the caudate toward visual value-guided behavior.

We next examined whether neural activity was differentially linked to behavior by correlating firing rates with subsequent finger-in RTs on a trial-by-trial basis. For bimodal value neurons, the proportions of neurons did not differ across the three striatal subregions in either T-VRT or V-VRT (Fig. 2I). Visual-selective value neurons also showed similar proportions across striatal subregions (Fig. 2J). The proportion of tactile-selective value neurons showed a marginal difference across striatal subregions in T-VRT. However, post hoc comparisons did not reveal significant differences between individual pairs of subregions, suggesting that this effect does not reflect clear differences between specific subregions.

Together, these results indicate that single-neuron properties across striatal subregions do not exhibit systematic differences and provide no evidence for a consistent organizational principle. Functional differences among striatal subregions are therefore unlikely to be captured by univariate features of neural activity, but may instead emerge from the structure of population-level representations.

### Neural geometry is differentially coupled to behavior across striatal subregions

To compare striatal subregions at the population level, we examined whether value representations converge to a shared representational geometry across blocks, and whether this convergence is enhanced during adapted value-guided behavior. This convergence can be assessed by cross-condition generalization performance (CCGP) ^24^, which measures how similar value representations remain across conditions. When neural representations are aligned such that value axes remain parallel across conditions, they enable a simple linear readout, allowing downstream circuits to efficiently decode value information for behavioral control ^25–29^. Thus, value generalization links population structure to value-guided behavior.

Consistent with this, recent studies suggest that generalized representations more closely track behavioral outputs than conventional decoding approaches ^24,30^. We therefore quantified CCGP separately for trials with adapted and unadapted value-guided behavior to determine whether value generalization across blocks reflects behavioral state across striatal subregions (Fig. S6A). Subregions contributing to sensory-specific value-guided behavior are expected to show greater CCGP modulation between adapted and unadapted trials.

Trials were classified into adapted and unadapted states based on RT patterns (Fig. 3A). ^30^. Adapted trials showed faster RTs for good objects and slower RTs for bad ones, while unadapted trials exhibited the opposite pattern (Fig. 3B). These results indicate that behavior was aligned with current value contingencies in adapted trials but misaligned in unadapted trials. CCGP was then compared between adapted and unadapted trials (Fig. 3C). For each subregion, CCGP in adapted trials was plotted against that in unadapted trials. Points falling below the identity line indicate higher CCGP during adapted than unadapted trials, reflecting stronger coupling between neural structure and behavior.

**Figure 3.**
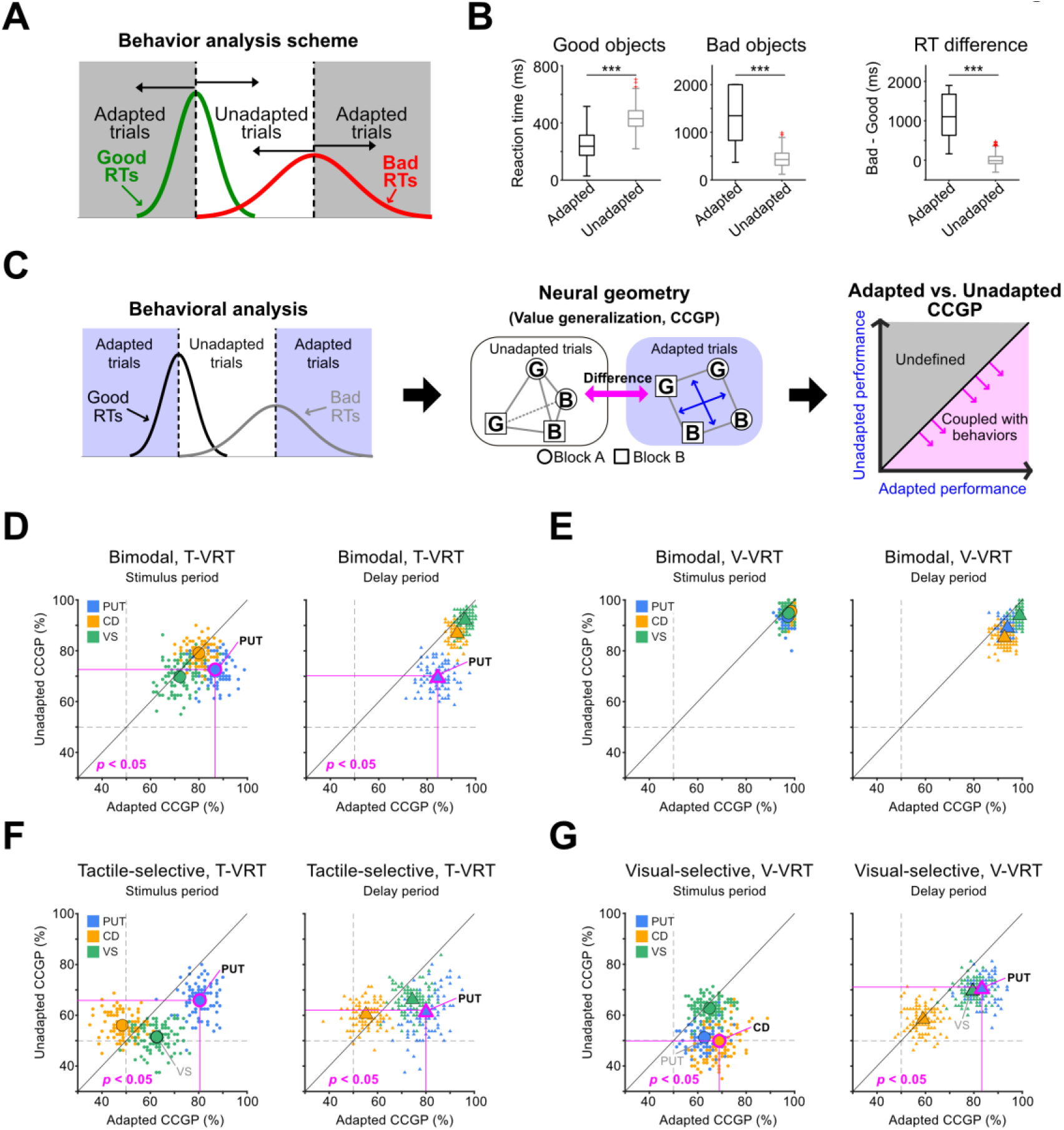
Neural geometry-behavior coupling difference across the striatal subregions. **(A)** Behavioral categorization scheme. Trials were classified into adapted and unadapted behavioral states based on the distribution of finger-in reaction times (RTs) for good and bad value stimuli. **(B)** Behavioral validation. Comparison of RTs for good stimuli, bad stimuli, and their difference (bad – good) between adapted and unadapted trials. Statistical details are provided in Table S2. **(C)** Analysis pipeline. Schematic illustrating the analytical workflow, from trial classification (adapted vs. unadapted) to CCGP computation and construction of the comparison plots. **(D-G)** CCGP comparison between behavioral states. **(D)** Bimodal value neurons in T-VRT. *Left*: stimulus period; *right*: delay period. Comparison plots show the relationship between adapted and unadapted CCGP performances. Colors denote recording regions: putamen (blue), caudate (yellow), and ventral striatum (green). Large symbols represent the mean across 100 iterations, whereas small colored symbols show individual iterations. Magenta outlines indicate performances significantly higher than chance (right-tailed z-test, *p* < 0.05). **(E)** Bimodal value neurons in V-VRT. Same conventions as in (D). **(F)** Tactile-selective value neurons in T-VRT. Same conventions as in (D). **(G)** Visual-selective value neurons in V-VRT. Same conventions as in (D). Box plot elements: center line, median; box limits, 25th and 75th percentiles; whiskers, 1.5 x IQR; red crosses, outliers. ***p < 0.0005; n.s., not significant.

In bimodal value neurons, the link between value generalization and behavior was most prominent in the putamen during T-VRT (Fig. 3D-E). The putamen neurons showed significantly higher CCGP differences between adapted and unadapted trials during both the stimulus and delay periods (Table S3), whereas the caudate and ventral striatum showed no significant difference across the behavioral state (Fig. 3D). Accordingly, CCGP modulation between adapted and unadapted trials was greatest in the putamen among subregions (Fig. S6B). This effect was absent in V-VRT, where all regions showed comparable CCGP regardless of behavioral state (Fig. 3E).

Similar results were found in tactile-selective value neurons. Tactile-selective value neurons in the putamen exhibited significantly greater CCGP differences than those in other striatal regions during both stimulus and delay periods (Fig. 3F). These results indicate that tactile value generalization is more strongly linked to behavior in bimodal and tactile-selective value neurons of the putamen than in other striatal subregions.

Visual-selective value neurons showed coupling between value generalization and behavior at distinct time points. In V-VRT, these neurons in the caudate showed significant coupling during stimulus presentation, whereas those in the putamen showed significant coupling during the delay period (Figs. 3G and S6C). Visual-selective value neurons in the ventral striatum showed a tendency toward coupling during the delay period in V-VRT (*p* = 0.097; Table S3). A similar pattern was observed for tactile-selective value neurons in the ventral striatum during the stimulus period of T-VRT (*p* = 0.065; Table S3) (Fig. 3F). Notably, these region-specific coupling patterns were uniquely revealed by CCGP but not by conventional SVM decoding (Fig. S6D-G) (Table S4).

Overall, our results indicate that neural population geometry and its coupling to behavior are not uniformly expressed across striatal subregions, but instead emerge selectively depending on the task. If this shared neural geometry underlies value-guided behavior, the causal influence of each striatal subregion should depend on the magnitude of difference in this geometry between adapted and unadapted trials (Table S3). Based on our T-VRT results, the putamen may mainly support tactile value-guided behavior, with additional contribution from the ventral striatum. In V-VRT, the caudate may support visual value-guided behavior during stimulus presentation, whereas the putamen and ventral striatum may contribute during the delay period. Yet, the causal role of neural geometry–behavior coupling in value-guided behavior remains unclear.

### Dissociable contributions of striatal subregions to modality-specific value-guided decisions

Based on these differences in neural geometry-behavior coupling across striatal subregions, we hypothesized that distinct subregions are causally required for modality-specific value-guided behavior. To test this, we performed reversible inactivation using the GABA_A_ receptor agonist muscimol while monkeys performed value-guided decision-making (Fig. 4A). Muscimol and saline (control) were injected into striatal subregions in both monkeys (Fig. S7A-B). Successful inactivation was confirmed by voltage traces showing reduced neural activity under muscimol compared to saline (Fig. S7C).

**Figure 4.**
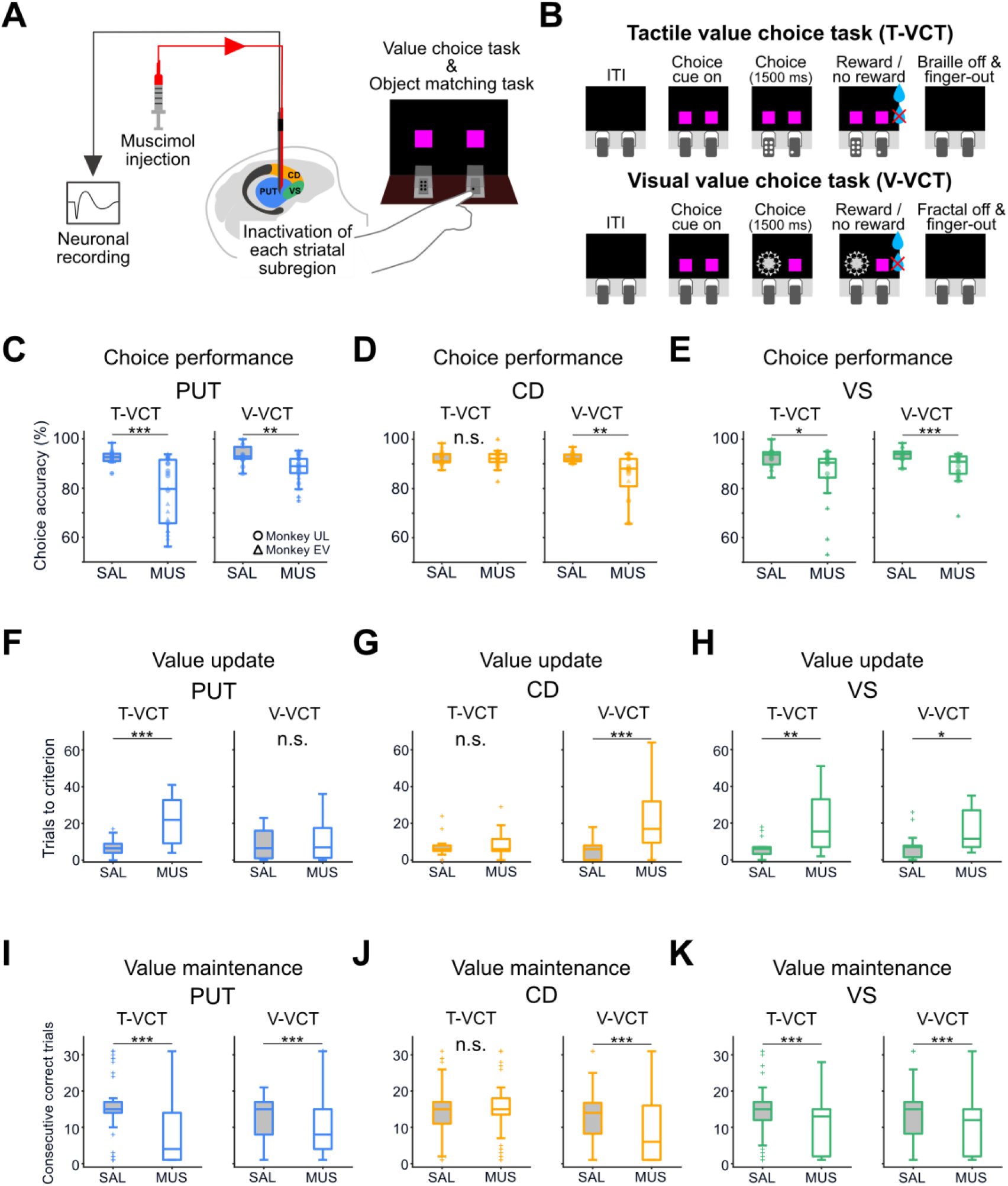
Causal contribution of striatal regions to value-guided choice behavior. **(A)** Experimental setup for reversible inactivation. Muscimol (or saline) was injected through a cannula-electrode system, enabling targeted perturbation and simultaneous recording of baseline activity to confirm inactivation. **(B)** Behavioral paradigms. Schematics of tactile (T-VCT) value choice task and visual (V-VCT) value choice task. The procedures were identical except for the stimulus modality. **(C-E)** Comparison of correct performance in T-VCT and V-VCT between saline and muscimol conditions in the putamen **(C)**, caudate **(D)**, and ventral striatum **(E)**. Statistical details are provided in Table S2. Symbols denote individual subjects (circle: Monkey UL; triangle: Monkey EV). **(F-H)** Analysis of value update, measured in trials-to-criterion in the putamen **(F)**, caudate **(G)**, and ventral striatum **(H)**. **(I-K)** Analysis of value maintenance, measured in consecutive correct trials, in the putamen **(I)**, caudate **(J)**, and ventral striatum **(K)**. Box plot elements: center line, median; box limits, 25th and 75th percentiles; whiskers, 1.5 x IQR; red crosses, outliers. *p < 0.05, **p < 0.005, ***p < 0.0005; n.s., not significant.

To determine how each striatal subregion contributes to sensory-driven decisions, we used value-based choice tasks in which monkeys chose between stimuli associated with different reward values (Figs. 4B and S7G-H). In both tactile (T-VCT) and visual (V-VCT) value choice tasks, value contingencies were reversed across blocks, requiring animals to update stimulus-reward associations to guide their choices (Figs. S7D-F and S8A).

We first assessed the effects of inactivation in each striatal subregion on task performance. Putamen inactivation significantly reduced the proportion of correct choices in both T-VCT and V-VCT, with a stronger and more consistent impairment in T-VCT (Fig. 4C). To quantify modality bias, we estimated the magnitude of impairment using the Hodges-Lehmann estimator, which revealed a larger performance decrement in T-VCT than in V-VCT (Fig. S8B, left panel). At the individual level, Monkey EV showed significant impairments in both tasks, with a greater deficit in T-VCT (Fig. S8F), whereas Monkey UL exhibited a significant deficit selectively in T-VCT, with no impairment in V-VCT (Fig. S8C). Thus, the putamen preferentially contributes to tactile value-guided decision-making compared to visual value-guided decision-making.

Caudate inactivation impaired choice accuracy only in V-VCT, with no effect on T-VCT (Fig. 4D). This was also confirmed by a larger performance decrement in V-VCT than in T-VCT, as quantified using the Hodges-Lehmann estimator (Fig. S8B, middle panel). Both monkeys exhibited significant deficits in V-VCT, with no impairment in T-VCT (Fig. S8D, G). These results indicate that the caudate is selectively engaged in visual value-guided decision-making with no clear contribution to tactile value-guided choices.

In contrast, ventral striatum inactivation impaired choice accuracy in both tasks (Fig. 4E), with comparable magnitudes of impairment across modalities (Fig. S8B, right panel). At the individual level, however, Monkey EV showed significant impairments in both tasks (Fig. S8H), whereas Monkey UL exhibited a significant impairment only in V-VCT (Fig. S8E). These results indicate that the ventral striatum contributes to value-guided decision-making, with variability in the strength of its effects across individuals. This variability may reflect differential contributions to value updating and maintenance processes, as subsequent analyses reveal.

To determine whether these effects reflect specific impairments in value-guided behavior, we examined performance in control tasks. Striatal inactivation did not impair sensory discrimination, working memory, or motor performance in control tactile and visual stimulus-matching tasks (Fig. S9). Trajectory analyses of eye and finger movements further confirmed that striatal inactivation did not affect these motor outputs (Fig. S10). These results indicate that the observed deficits are specific to value-guided decision-making.

### Dissociable causal roles of striatal subregions in sensory-specific value update and maintenance

Effective decision-making relies on the ability to flexibly update reward associations during reversal learning and to stably maintain them for consistent performance ^31–34^. Given that striatal subregions are involved in these processes ^35–40^, we sought to investigate their causal roles in value updating and maintenance across modalities.

To first examine the role of each subregion in value updating, we calculated trials-to-criterion, a metric reflecting how rapidly reward associations are updated. It was defined as the number of trials required for each monkey to reach a predefined number of consecutive correct choices (14 trials for Monkey UL and 5 trials for Monkey EV) (Fig. S11A).

Interestingly, we found that the putamen and caudate play distinct roles in value updating across sensory modalities. Putamen inactivation selectively impaired tactile value updating but not visual value updating (Fig. 4F): the number of trials-to-criterion significantly increased only in T-VCT, while remaining unchanged in V-VCT in both monkeys (Fig. S11C, F). In contrast, caudate inactivation selectively disrupted visual value updating but not tactile value updating (Fig. 4G). Trials-to-criterion significantly increased in V-VCT, whereas no effect was observed in T-VCT in both monkeys (Fig. S11D, G).

Conversely, the ventral striatum contributed to value updating across both modalities in both monkeys (Figs. 4H and S11E, H). These results show that the putamen selectively supports tactile value updating, the caudate in visual, and the ventral striatum contributes to value updating across modalities.

Next, we examined whether each striatal region contributes to maintaining value information that supports tactile or visual value-guided choices. To quantify this, we compared the number of consecutive correct trials between muscimol and saline conditions (Fig. S11B).

Putamen inactivation reduced consecutive correct trials in both tactile and visual tasks (Fig. 4I), although this effect was not consistent across individuals. Monkey EV showed significant deficits in both tasks, whereas Monkey UL exhibited a significant effect in T-VCT and a tendency in V-VCT (Fig. S11I, L). Ventral striatum inactivation impaired the maintenance of both tactile and visual values (Fig. 4K), and this effect was consistent across individuals (Fig. S11K, N). In contrast, caudate inactivation selectively impaired visual value maintenance in both monkeys (Figs. 4J and S11J, M).

Together, our inactivation results reveal that striatal subregions play distinct roles in tactile and visual value updating and maintenance. The caudate is specialized for visual value updating, the putamen for tactile value updating, and the ventral striatum contributes to both. In contrast, the putamen and ventral striatum preferentially support value maintenance across tactile and visual modalities, whereas the caudate selectively supports visual value maintenance.

## Discussion

Our results reveal functional dissociation across striatal subregions, with the putamen, caudate, and ventral striatum preferentially supporting tactile, visual, and integrative value-based decision-making, respectively. These differences were not explained by single-neuron activity or conventional decoding, but instead emerged from neural geometry–behavior coupling ^24,30,41^, indicating that behavioral relevance depends on how each subregion’s neural geometry is structured rather than how much information is encoded (Fig. S12).

Despite broadly similar value encoding across subregions, behavioral coupling differed, suggesting that functional dissociation arises from differences in readout accessibility. The caudate and putamen guide modality-specific decisions based on value, whereas the ventral striatum integrates across modalities, potentially serving a critic-like role and contributing to action selection ^42,43^. Under conflict, the ventral striatum integrates competing signals, whereas the caudate and putamen maintain modality-specific coding. These roles were most distinct during value learning and more convergent during maintenance, indicating differential contributions to updating versus stable expression of value ^37,44–46^. At the output level, value-guided decisions likely emerge from the weighted integration of parallel striatal signals at downstream structures such as the GPi, rather than from strictly segregated pathways^7,13,47–52^.

Together, our findings show that similar value signals can support distinct behavioral functions across striatal subregions. More broadly, they suggest a general principle by which similar neural representations distributed across brain regions are selectively recruited for behavioral control.

## Materials and Methods

### Braille presentation for the experiment

The apparatus used to present braille patterns to monkeys was the same as in our previous study (*23*). The braille presenter consisted of two opaque presentation cases into which monkeys inserted their index fingers to explore different braille patterns based solely on tactile input. A braille unit comprised six dots, and each dot was independently controlled by the experimenter according to the task schedule using the computer program BLIP, designed for electrophysiology and behavior studies for non-human primates (BLIP, Laboratory of Sensorimotor Research, National Eye Institute, National Institutes of Health, accessible at www.cocila.net/blip).

To precisely monitor finger entry and withdrawal from each presentation case, two photo interrupters (F249, Arduino module) were positioned at the entrance of each case. Two endoscopic cameras (PS-EC200) were also placed at the end of each braille presentation case, and an overhead camera (D455, Intel, USA) was installed above the apparatus for real-time monitoring and motion tracking analysis using DeepLabCut (version 2.2.2).

### Sensory stimuli

For visual stimuli, we used two fractal images that were used in our previous studies (*23*, *30*). For the inactivation experiments using muscimol, two black-and-white fractal images were used. Fractal images were generated using a MATLAB-based program (https://github.com/ProfKimHF/fractalgenerator). The fractal images were approximately 20° x 20° in size, while the squares were 8° x 8° in size.

For tactile stimuli, we used two different braille patterns from our previous studies (*23*, *30*). Each braille unit was configured such that each presentation case produced a single braille pattern. Each dot had a diameter of 2 mm and a height of 1.5 mm, and each braille pattern measured 10 x 5 mm.

### General procedures

Two rhesus macaques (Macaca mulatta; 4.5 kg female Monkey EV and 9.8 kg male Monkey UL) were used in this study. All experimental and animal care procedures were approved by the Institutional Animal Care and Use Committee of Seoul National University (SNU-251017-3). Under general anesthesia and aseptic surgical conditions, both monkeys were implanted with a plastic or titanium head holder and a plastic recording chamber. Each chamber was tilted laterally by 25° to align with the striatum (putamen, caudate, and ventral striatum). After the monkeys recovered from surgery (> 1 month), training and recording sessions commenced.

### Single-unit recording

While the monkeys were performing tasks, the activity of single neurons in the striatum (putamen, caudate, and ventral striatum) was recorded using conventional electrophysiological methods. We determined recording sites using a grid system with 1 mm spacing and MR images acquired with a 3T scanner (Siemens), aligned with the angle of the recording chamber. For the electrophysiology probe, we inserted a glass-coated electrode (Alpha-Omega) into the brain through a stainless-steel guide tube using an oil-driven micromanipulator (MO-974A, Narishige). Neuronal signals from the electrode were amplified, filtered (250 Hz–10 kHz), and digitized (at a 30 kHz sampling rate with 16-bit A/D resolution) through a Scout system (Ripple Neuro, UT). These neuronal signals were then isolated online using a custom voltage-time window discrimination software (BLIP), with event timings recorded at 1 kHz. Each spike waveform was sampled at 50 kHz. To minimize potential temporal confounds, we alternated recording sessions among the striatal subregions rather than completing recordings in one region before moving to another. This ensured that neuronal samples from each region were collected throughout the entire experimental period.

### Behavioral tasks for electrophysiology

Monkeys performed two sensory modality-specific value tasks to test neuronal activity in the striatum, as described in our previous study: the Tactile Value Reversal Task (T-VRT) and the Visual Value Reversal Task (V-VRT) (*23*).

In the T-VRT single-stimulus trial (Fig. S1A, C), monkeys were required to insert their left index finger into the left hole of the braille presenter after the onset of the first finger-in cue. After finger insertion, one of two braille patterns was then delivered and remained for 500 ms. The holes presenting braille patterns were not visible to the monkeys, so they had to rely on tactile input to discriminate and learn the value of the stimuli. After both the braille pattern and the finger-in cue had disappeared, the monkeys were instructed to withdraw their finger. A delay period of 500–1000 ms followed, during which no stimuli were presented. After the delay, the second finger-in cue appeared, and the monkeys reinserted their finger to touch the same stimulus presented during the first finger insertion and received a stimulus-associated reward, or they could withhold reinserting by waiting 1000 or 2000 ms. After choosing to reinsert their finger, they had to wait 200 ms before reward delivery.

Following four single-stimulus trials, a choice trial was presented. Each choice trial began with the simultaneous appearance of two magenta dots on the monitor, prompting the monkey to insert its index finger into one of the two holes. During this phase, monkeys were permitted to freely explore both holes to sample the tactile (braille) stimuli. A final selection was registered when the monkey maintained its finger within a single hole for 1300–1500 ms.

The finger insertion reaction time was measured from the onset of the second finger­in cue to finger entry into the hole in the single-stimulus trial. In each session, two blocks were included, each consisting of 50 trials. In each block, the stimulus-reward association was reversed. Blocks were changed without notification, requiring the monkeys to learn the new reward contingency by trial and error. The order of the blocks was randomized across sessions, ensuring that monkeys were unaware of block identity until the start of each session.

The procedure of V-VRT (Fig. S1B, D) was the same as that of T-VRT, except for the sensory stimuli associated with rewards. Instead of braille patterns, fractal images were displayed on a monitor in front of the monkeys as they inserted their finger into the hole. V-VRT and T-VRT were conducted separately. For Monkey UL, magenta and blue cues were used (Fig. 1A, B), and for Monkey EV, white cues were used instead of colored cues (Fig. S1C, D).

### Behavioral tasks for muscimol inactivation study

In the muscimol inactivation study, the monkeys performed both value and control tasks. For the value task, they performed the Tactile Value Choice Task (T-VCT) and the Visual Value Choice Task (V-VCT), which measured how well the monkeys learned and associated tactile or visual stimuli with reward value. For the control task assessing basic sensory, motor, and working memory demands, the Tactile Stimulus Matching Task (T-SMT) and the Visual Stimulus Matching Task (V-SMT) were performed to examine the monkeys’ ability to discriminate braille patterns and fractal images, respectively.

For Monkey UL, in T-VCT, after the onset of two finger-in cues, the monkey could insert its finger into either hole and explore which stimulus was presented in each hole (Fig. 4B). The exploration period lasted as long as the monkey chose. The monkey chose one stimulus by inserting its finger into one hole and maintaining the insertion for more than 1500 ms. After the choice, the reward associated with the chosen stimulus was delivered 200 ms after cue and stimulus offset. The reward-contingency schedule was also different between monkeys. For Monkey UL, each session consisted of 100 trials, and blocks were reversed after the monkey made 14–19 consecutive correct trials (Fig. S7D).

For Monkey EV, the sequence of T-VCT was slightly different because simultaneous presentation of two cues at the beginning of a trial interfered with stable task performance; therefore, the task was modified to match the initial trial sequence of T-VRT. (Fig. S7G, H). Each session consisted of repeated cycles of one single-stimulus trial that only showed the good stimulus and four choice trials (Fig. S7E). In each session, there were 80 trials with two blocks: Block A and B. The order of the blocks was unknown until the start of the session (Fig. S7F). The first part of each choice trial was the same as that in T-VRT: the monkey was required to insert its finger into the left hole after the onset of the first finger-in cue, and after finger withdrawal following cue and stimulus offset, a choice trial with two cues began. The location of each stimulus remained identical between the first finger insertion and the choice phase. In other words, during the first finger insertion, the monkey sampled the stimulus in the left hole and retained this information to guide the subsequent choice phase.

### Classification of single-neuron responses

To classify different types of value neurons, we used the following procedure, as previously described (*23*, *30*). First, to isolate neurons relevant to the behavioral task, we screened for units that modulated their firing rates significantly during task events. We defined a baseline control window as the 200 ms interval preceding the first finger-in cue. This baseline activity was compared against neural activity within a 500 ms test window starting from the onset of ten specific task events: (1) first cue onset, (2) first finger insertion, (3) stimulus onset, (4) stimulus offset, (5) first finger withdrawal, (6) second cue onset, (7) second finger insertion, (8) reward delivery, (9) second cue offset, and (10) second finger withdrawal. For each event, a Wilcoxon rank-sum test was performed across trials, and neurons showing significant modulation during at least one event epoch were classified as task-related neurons.

Next, within this task-related population, we investigated whether neurons encoded value information. Spike counts were analyzed during the stimulus presentation period (0–500 ms after stimulus onset) and the subsequent delay period (0–500 ms after finger withdrawal). To determine value selectivity, firing rates under ‘good’ and ‘bad’ value conditions were compared using a Wilcoxon rank-sum test.

Finally, based on the sensory modality through which these value signals appeared, neurons were categorized into three functional groups. Neurons that discriminated value significantly only during T-VRT (but not V-VRT) were defined as tactile-selective value neurons. Conversely, those showing significant value discrimination only during V-VRT were labeled visual-selective value neurons. Neurons that encoded value in both tasks were classified as bimodal value neurons. A detailed summary of these neuronal counts and definitions is provided in Supplementary Table S1.

### Value discrimination latency and duration

To precisely quantify the temporal evolution of value coding—specifically when the signal emerged and how long it persisted—we analyzed the firing rate dynamics of value neurons for each sensory modality (*30*). Instead of relying on predefined fixed epochs, we employed a sliding window approach to capture fine temporal dynamics. We calculated spike counts within a 100 ms moving window, shifting the window by 1 ms increments across the trial duration.

At each time step, we assessed whether each neuron distinguished between good and bad stimuli by comparing spike counts using a Wilcoxon rank-sum test. To control for the false discovery rate (FDR) across multiple time points, we adjusted the raw p-values using the Benjamini–Hochberg procedure. Based on these corrected p-values, we derived two key temporal metrics:

1. Value onset latency: To determine the signal’s initiation, we identified the first time point after stimulus onset where the neural activity showed significant differentiation for a duration of at least 10 consecutive bins (10 ms).
2. Value discrimination duration: To measure the total persistence of the value signal, we calculated the sum of all time bins that exhibited statistically significant value discrimination, regardless of their continuity.

Finally, to evaluate whether these temporal features differed by modality, we compared the distributions of onset latencies and durations across striatal regions. Statistical significance was assessed using a Kruskal–Wallis test (*p* < 0.05).

### Correlation between neural response and finger-in reaction time

We sought to determine whether trial-by-trial fluctuations in neural firing rates predicted the speed of the monkeys’ finger insertion (reaction time, RT). We calculated spike counts within specific 500 ms time windows tailored to neuronal subtype: relative to stimulus onset (0–500 ms) for stimulus value neurons, and relative to finger withdrawal (0–500 ms) for delay value neurons. To capture the dynamic evolution of neural responses and behavior, particularly during value reversals, we grouped trials into bins of two. Because the task design involved pseudo-randomized presentation of good and bad stimuli, these bins were organized based on the cumulative number of presentations for each stimulus type within a block, ensuring a temporally ordered sequence of learning. We then performed Pearson’s correlation analysis to quantify the linear relationship between spike counts and RTs for each neuron.

### Value magnitude index

To quantify the strength of value discrimination, we calculated the Receiver Operating Characteristic (ROC) curve for each neuron. The analysis compared the distribution of spike counts between good- and bad-value conditions in both T-VRT and V-VRT tasks. The area under the ROC curve served as a quantitative measure of discrimination strength. An ROC value of 0.5 implies that the neuron responds identically to both conditions (no selectivity), whereas values approaching 1.0 or 0.0 indicate strong selectivity for one value (good or bad) over the other. Specifically, ROC values > 0.5 denote positive value coding (preferring good), while ROC values < 0.5 denote negative value coding (preferring bad).

### Population decoding analysis

To prepare the data for population analysis, we first isolated the neural activity relevant to value processing. Since our initial analysis indicated that the striatum encodes value during two distinct phases—stimulus presentation and the subsequent delay period—we extracted spike trains specifically from these periods. We defined the stimulus window as 0 to 500 ms aligned to stimulus onset, and the delay window as 0 to 500 ms aligned to finger withdrawal. We then concatenated these two segments into a single time series for analysis. This selective concatenation, rather than using a continuous time window from stimulus onset to the end of the delay, was designed to exclude neural activity related to the finger movement, ensuring that decoding results primarily reflected value processing rather than motor activity. To prevent neurons with naturally high firing rates from dominating the analysis, we normalized the concatenated spike trains using z-score normalization, capturing relative response changes across the entire population.

For the decoding analysis, spike counts were calculated during the stimulus and delay periods. Using these pre-processed neural activities, we constructed pseudo-populations by pooling neurons and employed a linear Support Vector Machine (SVM) decoder with 10-fold cross-validation. We repeated this process for 100 iterations, generating random pseudo-populations for each run, and reported the mean decoding accuracy across iterations. Throughout this process, we strictly balanced the data by including an equal number of trials, with at least 10 trials for each condition (block x value combinations), and the same number of neurons across the three striatal regions for each analysis (bimodal, n = 83; tactile-selective, n = 24; visual-selective, n = 34). Finally, data were pooled across both monkeys, as the key neural features were consistent between the two subjects.

### Cross-condition generalization performance (CCGP)

We utilized CCGP to quantify the generalization capability of neural populations across shared task variables (*30*, *31*). In contrast to traditional decoding methods, in which training and test sets share condition identities, CCGP requires the decoder to predict labels for conditions not seen during training. This cross-condition validation assesses whether neural geometry preserves a consistent parallel structure across shared variables. To decode the generalizability of value (good vs. bad) across different task blocks (Block A vs. Block B), we computed CCGP across all four possible training–testing permutations based on the four condition combinations. Specifically, a linear decoder was trained and tested on the following four data splits: (1) train on Block A (good vs. bad) and test on Block B (good vs. bad); (2) train on Block B and test on Block A; (3) train on a mixed-block subset (Block A-good vs. Block B-bad) and test on the remaining subset (Block B-good vs. Block A-bad); and (4) train on the alternate mixed subset (Block B-good vs. Block A-bad) and test on the remaining subset (Block A-good vs. Block B-bad). The decoding results were averaged over 100 iterations for each permutation, and the mean across all four permutations was reported as the final generalization metric.

### Permutation model for testing significance of decoding performance

To verify that our SVM and CCGP performances were significantly above chance, we implemented a permutation test as described previously (*24*, *30*). We generated a null distribution by randomly shuffling the condition labels for each neuron while preserving the trial structure. This shuffling process was repeated 1,000 times for every analysis window, allowing us to construct a null distribution of decoding accuracies expected under the null hypothesis. We visualized these benchmarks by plotting the theoretical chance level in the main figures and the 95% confidence intervals (5th–95th percentiles) of the null distribution in the box plots. Statistical significance was then assessed by comparing the actual decoding performance with this null distribution using a right-tailed z-test, with p < 0.05 considered significant.

### Pharmacological inactivation of striatal regions

To investigate the causal contributions of specific striatal subregions to value processing, we reversibly inactivated the anterior putamen, caudate, and ventral striatum using the GABAA agonist muscimol. As a control, sterile saline was injected into the same regions in separate sessions. To ensure precise localization, we employed a custom-made injectrode system, consisting of an epoxy-coated tungsten microelectrode integrated with a silica injection tube (*36*). This setup allowed us to monitor neuronal activity immediately before and during injection. Successful inactivation was further validated by comparing voltage traces between muscimol and saline conditions, confirming suppression of neural activity (Fig. S7C).

Inactivation experiments were conducted after the completion of all baseline behavioral and neuronal data collection. Due to the large volume of the dorsal striatum, muscimol was injected at both dorsal and ventral locations within the putamen or caudate during a single session to ensure sufficient coverage. The ventral striatum received a single injection. At each site, we delivered 0.8–2.0 μl of muscimol (5.12 mM, Sigma) or saline at a controlled rate of 0.2 μl/min. Following a 10 min diffusion period, the monkeys began the behavioral tasks. The tasks were presented in a randomized order, and data collection continued for 2–3 h post-injection. Behavioral performance was compared between muscimol and saline sessions to quantify the effects of inactivation.

To minimize potential confounding factors such as order effects or long-term plasticity, the order of injection sites and injected substances was counterbalanced across sessions. Experiments were conducted weekly to allow for full washout, typically alternating between muscimol and saline treatments (e.g., muscimol in week *N*, saline in week *N+1*). The total number of sessions was balanced across conditions: Monkey UL received muscimol 4 times per region, with saline controls performed 3 times in the putamen, 4 in the caudate, and 3 in the ventral striatum (Fig. S6A, B). Monkey EV received muscimol 3 times in the putamen and 2 times each in the caudate and ventral striatum, with 2 saline control sessions per region (Fig. S6A, B).

### Behavioral analysis of value updating and maintenance

To investigate the contribution of striatal subregions to value updating and maintenance across two sensory modalities, we quantified trials-to-criterion and consecutive correct trials, respectively. For value updating analysis, we measured the number of trials (including both correct and incorrect responses) required for a subject to achieve a predefined number of consecutive correct choices. This criterion was set at 14 consecutive correct trials for Monkey UL and 5 for Monkey EV. Notably, the trials-to-criterion count included all behavioral attempts preceding the criterion, while excluding the criterion trials themselves.

To evaluate value maintenance, we counted all consecutive correct trials and compared these between muscimol and saline conditions. For this analysis, we excluded the initial trial of every block following a value reversal.

### Analysis of eye movements

Eye positions were continuously monitored at a 1 kHz sampling rate using high-speed infrared camera systems. Cameras were Oculomatic Pro (Open Ephys, USA) and ISCAN (ISCAN, USA). To assess the similarity between eye trajectories across reward value and experimental conditions, we utilized Dynamic Time Warping (DTW). This algorithm aligns two temporal sequences by identifying an optimal path that minimizes the total accumulated distance between them. Each trajectory was treated as a two-dimensional time series consisting of horizontal and vertical coordinates. The divergence between trajectories was determined by calculating the Euclidean distance between corresponding time points and minimizing the overall cost of alignment. Lower DTW distances indicated higher similarity, while higher values represented greater differences in movement patterns.

Furthermore, we used a linear decoding framework to determine if eye movement patterns significantly differed between experimental categories. The decoder was trained and tested to distinguish between within-condition distances and between-condition distances. To ensure the stability of the decoding performance, this procedure was conducted over 100 iterations using random sampling. The statistical significance of the decoder’s accuracy was validated against a null distribution generated through 1,000 permutations with shuffled labels, followed by a right-tailed z-test.

### Analysis of finger movements

We tracked the index finger’s motion during the experimental trials using the DeepLabCut (version 2.2.2) toolbox in conjunction with MATLAB (MathWorks, USA). High-speed video recordings of the finger were synchronized with neural and behavioral data through the Scout system and Trellis software (Ripple Neuro, USA). Specifically, we labeled three distinct points on the finger manually for 336 frames taken from six videos for each monkey (of which 95% were used for training). We used a ResNet-50-based neural network with default parameters for 3 number of training iterations. We validated with 1 shuffle, and found that the test error was: 6.7 (EV_left), 15.08 (UL_left) pixels, whereas the training error was: 3.25 (EV_left), 3.36 (UL_left) pixels (image size was 640 x 480 pixels). We then used a p-cutoff of 0.6 to filter the x-y coordinates for subsequent analyses. This network was then used to analyze videos from similar experimental settings. For each frame, the spatial position of the finger was determined by calculating the center of gravity of these three labeled points. To compare the resulting movement trajectories across different experimental conditions, we applied DTW, which measures the similarity between time-series data by finding an optimal alignment that minimizes the total distance between paths.

Using the same decoding procedure as for eye movements, a linear decoder was employed to evaluate whether the finger’s physical trajectories contained enough information to distinguish between different values and experimental conditions. The classification performance was validated using a repeated random sampling procedure and compared against a null distribution generated through 1,000 permutations to establish statistical significance.

### Data analysis tool and statistics

All behavioral and neuronal data analyses were performed using MATLAB (R2024b). A comprehensive summary of all statistical results, including test statistics and p-values, is provided in Table S2.

## Funding

This work was supported by the Bio&Medical Technology Development program (RS-2025-02263832), the Basic Science Research Program (RS-2024-00339355), the Global-LAMP program (RS-2023-00301976), and the ASTRA program (RS-2024-00436783) through the National Research Foundation (NRF) of Korea. We thank D.I. Ko for technical support and members in Institute of Laboratory Animal Resources (ILAR), SNU for technical assistance.

## Author contributions

H.F.K. supervised the entire project. JW. L. and H.F.K. designed the experiments. JW. L., MS. K., and SH. H. performed the behavior, single-unit recording, and inactivation experiments. JW. L. analyzed the data and prepared the figures. JW. L., SH. H., MS. K. and H.F.K wrote the first draft, and JW. L., SH. H., MS. K., and H.F.K. interpreted data and wrote the final manuscript.

## Competing interests

The authors declare no competing interests.

## Data availability

Due to ethical and regulatory restrictions, the dataset supporting this study cannot be publicly archived. Access to the data requires prior approval from the Institutional Animal Care and Use Committee of Seoul National University and National Research Foundation of Korea. Requests will be reviewed within several months and, if approved, data will be made available for academic, non-commercial use. Source data are provided with this paper.

## Code availability

Custom code associated with this study is available at https://github.com/ProfKimHF.

## Supplementary Information

**Supplementary Figure 1.**
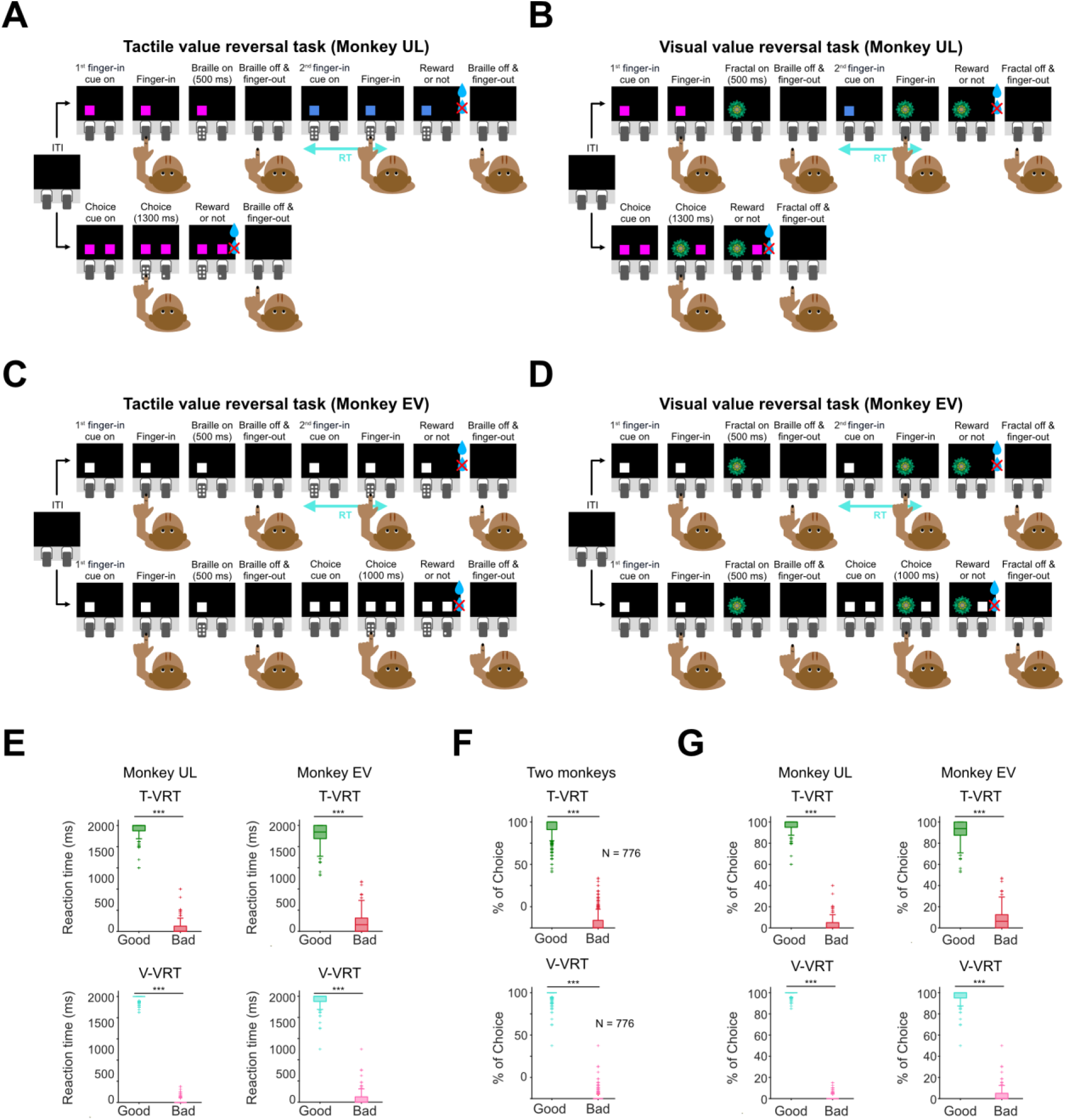
Task design and behavioral performance in tactile and visual value reversal tasks. **(A-B)** Full task scheme for Monkey UL. **(A)** Tactile value reversal task (T-VRT) with choice trials and **(B)** visual value reversal task (V-VRT) with choice trials. **(C-D)** Task modifications for Monkey EV. Schematics of the **(C)** T-VRT and **(D)** V-VRT adapted for Monkey EV. White cues were employed for Monkey EV. **(E)** Reaction times to good and bad stimuli in both monkeys. Comparison of finger-insertion RT differences (bad – good) in T-VRT (top) and V-VRT (bottom). *Left*: Reaction time in Monkey UL (n = 482, two-tailed paired t-test, p < 0.0001 for both T-VRT and V-VRT). *Right:* Monkey EV (n = 294, two-tailed paired t-test, p < 0.0001 for both T-VRT and V-VRT). **(F)** Percentages of good and bad choices in both monkeys (n = 776, two-tailed paired t-test, p < 0.0001 for both T-VRT and V-VRT). **(G)** Choice performance in individual monkeys. *Left*: Monkey UL (n = 482, two-tailed paired t-test, p < 0.0001 for both T-VRT and V-VRT). *Right*: Monkey EV (n = 294, two-tailed paired t-test, p < 0.0001 for both T-VRT and V-VRT). Box plot elements: center line, median; box limits, 25th and 75th percentiles; whiskers, 1.5 x IQR; red crosses, outliers. \**p* < 0.05, \*\**p* < 0.005, \*\*\**p* < 0.0005; n.s., not significant.

**Supplementary Figure 2.**
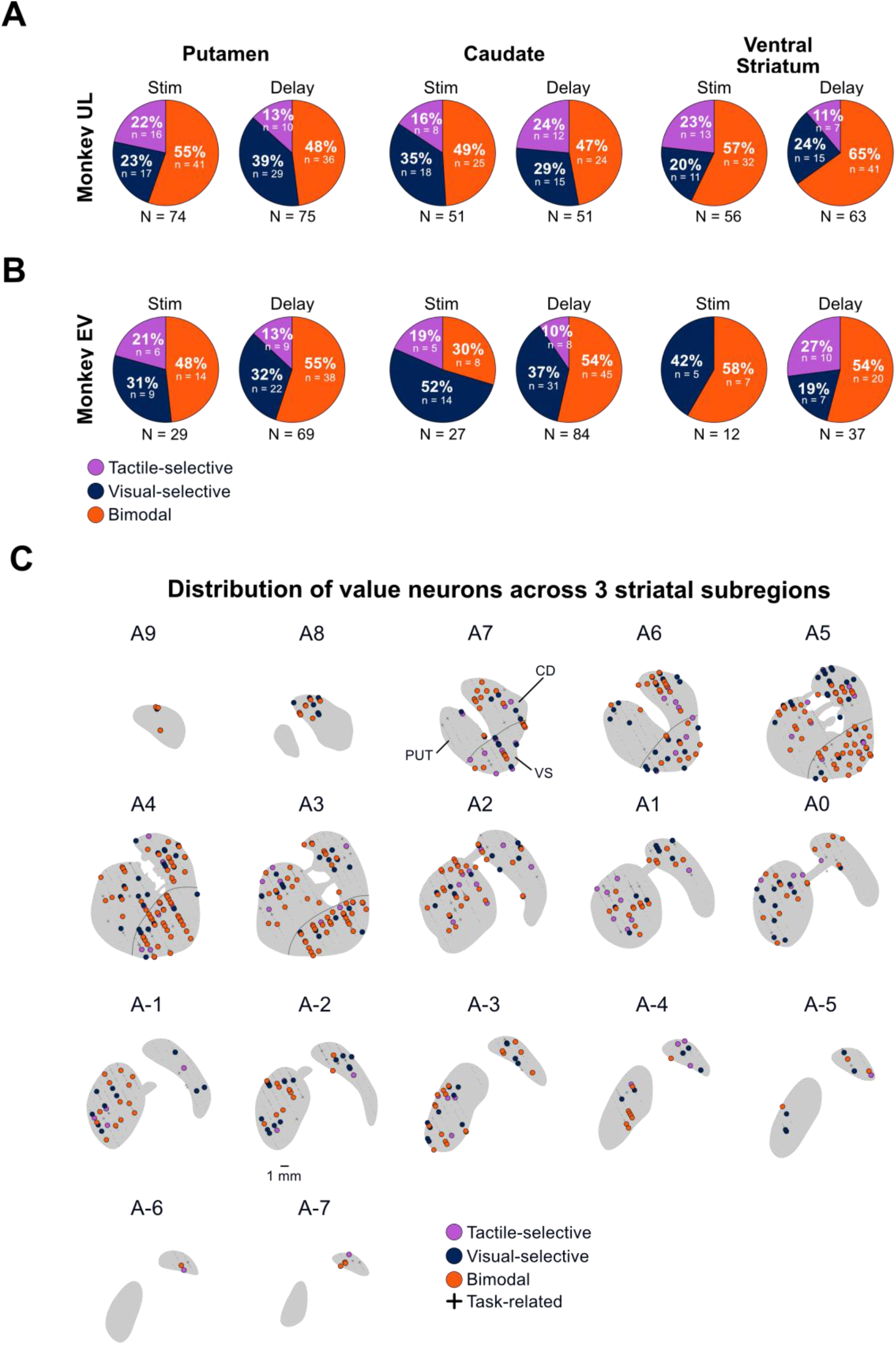
Subject-wise distribution of value neurons and reconstruction of recording sites. **(A-B)** Proportions of value neuron types in each subject, classified according to encoding epochs. **(A)** Results for Monkey UL. **(B)** Results for Monkey EV. **(C)** Reconstruction of recording sites. MR-based reconstruction of recording sites plotted on coronal sections of the striatum, including the putamen, caudate, and ventral striatum. Numbers indicate the anterior-posterior distance (in mm) from the anterior commissure (AC). Small black dots represent all neurons encountered during recording sessions, while crosses denote non-value neurons. Value neurons are color-coded: tactile-selective (purple), visual-selective (navy), and bimodal (orange).

**Supplementary Figure 3.**
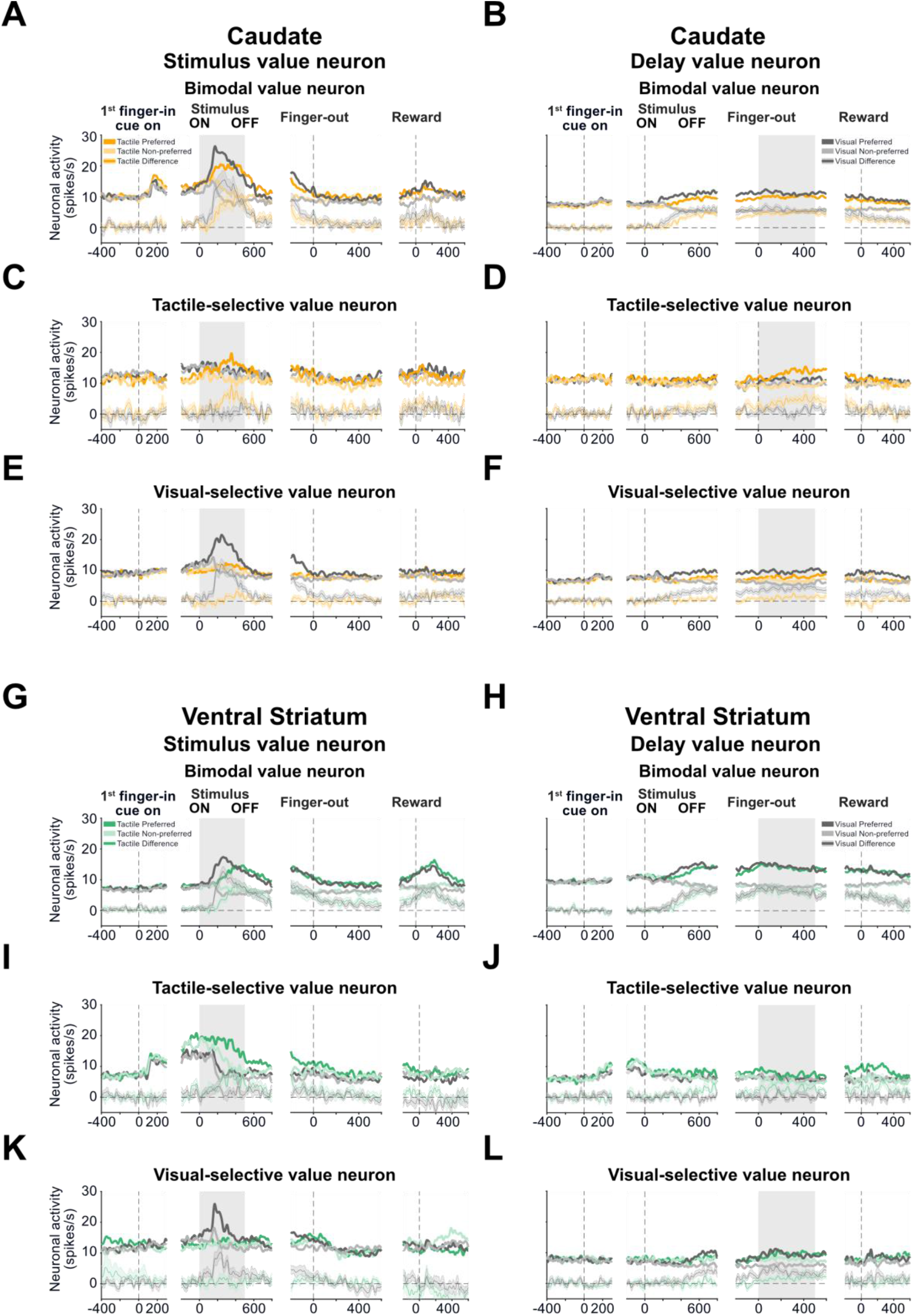
Stimulus and delay responses of value neurons in the caudate and ventral striatum. **(A–F)** Neural responses in the caudate. Averaged neural activity of the three types of value neurons during the stimulus and delay periods. **(A, B)** Bimodal value neurons (stimulus, *n* = 33; delay, *n* = 69), **(C, D)** tactile-selective value neurons (stimulus, *n* = 13; delay, *n* = 20), and **(E, F)** visual-selective value neurons (stimulus, *n* = 32; delay, *n* = 46). **(G-L)** Neural responses in the ventral striatum. Same conventions as in **(A-F)**. **(G, H)** Bimodal value neurons (stimulus, *n* = 39; delay, *n* = 61), **(I, J)** tactile-selective value neurons (stimulus, *n* = 13; delay, *n* = 17), and **(K, L)** visual-selective value neurons (stimulus, *n* = 16; delay, *n* = 22).

**Supplementary Figure 4.**
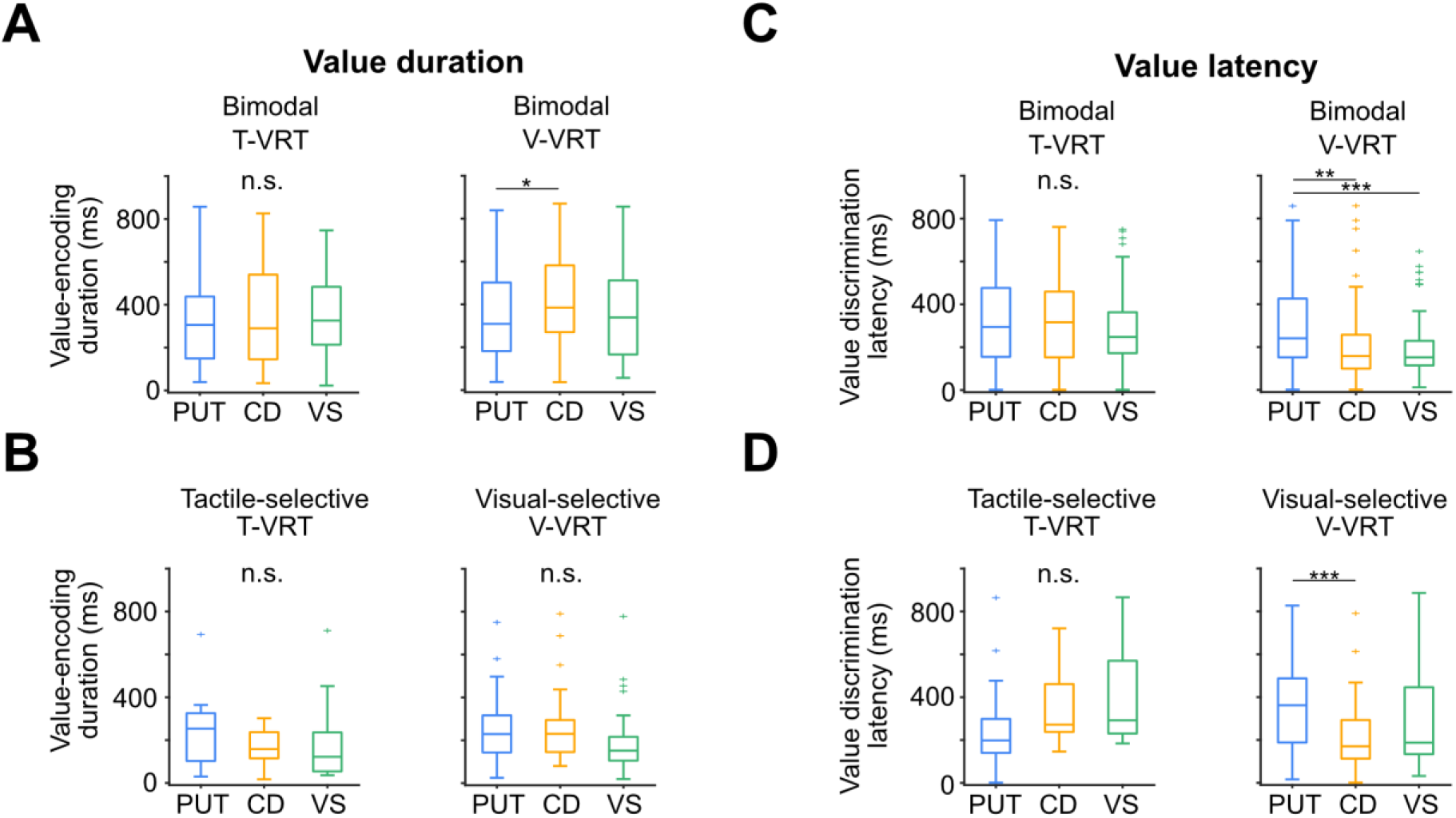
Single-cell characteristics of value encoding. **(A)** Value-encoding duration of bimodal value neurons. Bimodal value neurons during T-VRT (Kruskal-Wallis test *H*(2) = 2.60, p = 0.2727) and V-VRT (*H*(2) = 6.25, p = 0.0439, Bonferroni post hoc: putamen vs. caudate, p = 0.0468). **(B)** Value-encoding duration of modality-selective value neurons. Tactile-selective value neurons during T-VRT (Kruskal-Wallis test *H*(2) = 2.86, p = 0.2397), and visual-selective value neurons during V-VRT (*H*(2) = 3.93, p = 0.1401). **(C)** Value discrimination latency of bimodal value neurons. Bimodal value neurons during T-VRT (Kruskal-Wallis test *H*(2) = 1.33, p = 0.5140) and V-VRT (*H*(2) = 14.54, p = 0.000695, Bonferroni post hoc: putamen vs. caudate, p = 0.0101; putamen vs. ventral striatum, p = 0.0015). **(D)** Value discrimination latency of modality-selective value neurons. Tactile-selective value neurons during T-VRT (Kruskal-Wallis test H(2) = 5.70, p = 0.0577), and visual-selective value neurons during V-VRT (*H*(2) = 12.97, p = 0.0015, Bonferroni post hoc: putamen vs. caudate, *p* = 0.000983). Box plot elements: center line, median; box limits, 25th and 75th percentiles; whiskers, 1.5 x IQR; red crosses, outliers. \**p* < 0.05, \*\**p* < 0.005, \*\*\**p* < 0.0005, n.s., not significant.

**Supplementary Figure 5.**
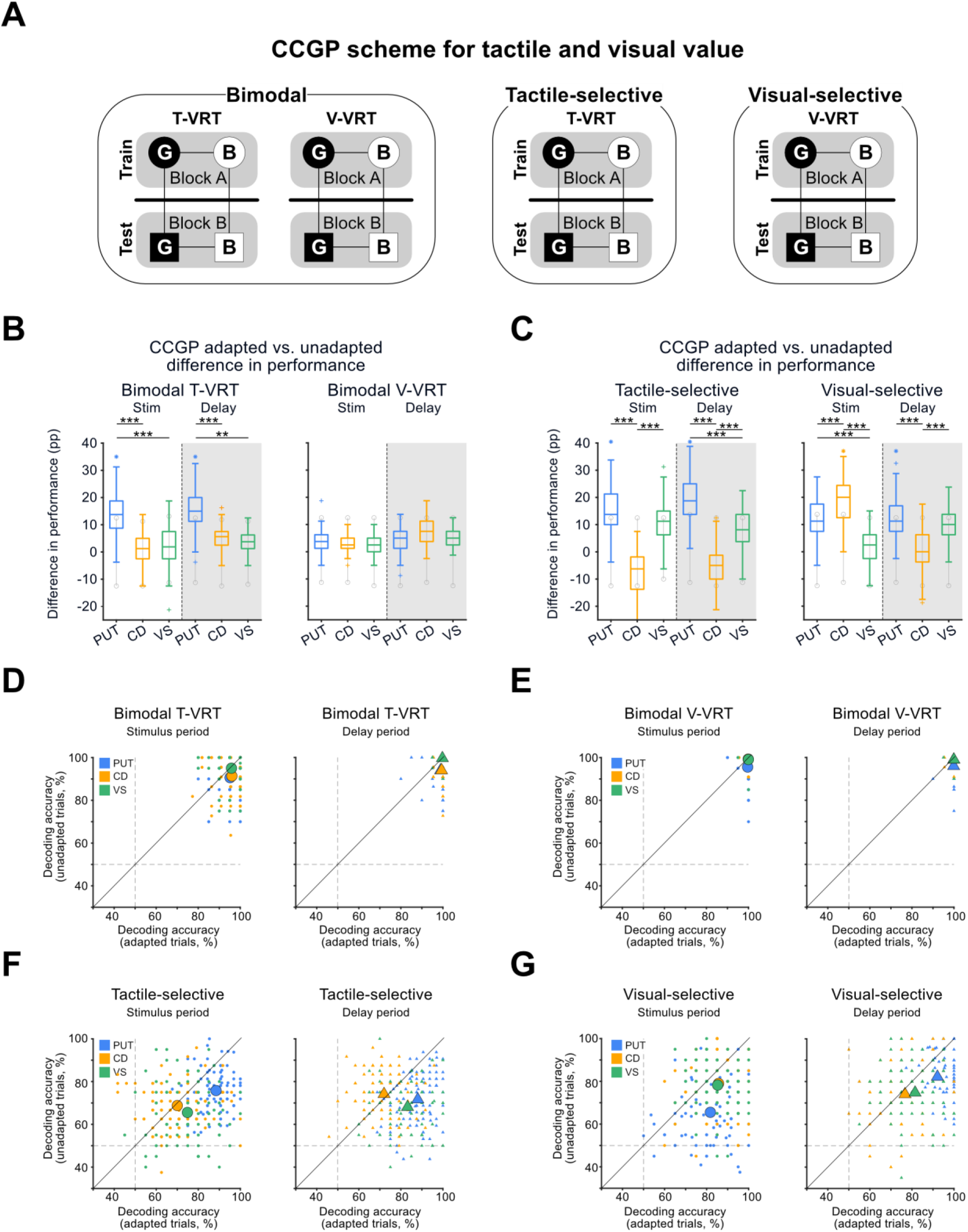
CCGP and SVM decoding of adapted and unadapted trials. **(A)** Schematic of Cross-condition generalization performance (CCGP) analysis. Value generalization was tested across different blocks (conditions) to evaluate the abstract representation of value. Circles and squares denote Block A and Block B, respectively, while black and white fills represent good and bad values. A decoder trained on data from one block was tested on the other to measure generalization capability. This block-wise approach allowed CCGP analysis of modality-selective neurons. **(B-C)** CCGP differences in adapted and unadapted trials. Colored asterisks indicate performance significantly exceeding chance levels (permutation test; right-tailed z-test, *p* < 0.05). Gray vertical lines with open circles represent the 95% confidence interval of the permutation-based null model. **(B)** CCGP differences in bimodal value neurons during T-VRT (*left panel*: One-way ANOVA, *F*(5, 594) = 109.75, *p* < 0.0001, post hoc tests, stimulus putamen vs. caudate, *p* < 0.0001; stimulus putamen vs. ventral striatum, *p* < 0.0001; delay putamen vs. caudate, *p* < 0.0001; delay putamen vs. ventral striatum, *p* < 0.0001) and V-VRT (*right panel*: Performances did not significantly differ from chance levels, and no inter-regional differences were tested). **(C)** CCGP differences in tactile-selective value neurons during T-VRT (*left panel*: Kruskal-Wallis test, *H*(5) = 389.25, *p* < 0.0001; post hoc tests, stimulus putamen vs. caudate, *p* < 0.0001; stimulus caudate vs. ventral striatum, *p* < 0.0001; delay putamen vs. caudate, *p* < 0.0001; delay putamen vs. ventral striatum, *p* < 0.0001; delay caudate vs. ventral striatum, *p <* 0.0001), and in visual-selective value neurons during V-VRT (*right panel*: Kruskal-Wallis test, *H*(5) = 280.40, *p* < 0.0001; post hoc tests, stimulus putamen vs. caudate, *p* < 0.0001; stimulus putamen vs. ventral striatum, *p* < 0.0001; stimulus caudate vs. ventral striatum, *p* < 0.0001; delay putamen vs. caudate, *p* < 0.0001; delay caudate vs. ventral striatum, *p <* 0.0001). **(D-G)** SVM analysis across cognitive states. **(D)** Comparison of decoding accuracies between adapted and unadapted behavioral trials during the stimulus and delay periods for bimodal value neurons in T-VRT. The x- and y-axes represent decoding accuracies in adapted and unadapted trials, respectively. Colors denote recording regions: putamen (blue), caudate (yellow), and ventral striatum (green). Large symbols represent the mean across 100 iterations, whereas small colored symbols represent individual iterations. **(E)** Bimodal value neurons in V-VRT. Same conventions as in (D). **(F)** Tactile-selective value neurons in T-VRT. Same conventions as in (D). **(G)** Visual-selective value neurons in V-VRT. Same conventions as in (D). Box plot elements: center line, median; box limits, 25th and 75th percentiles; whiskers, 1.5 x IQR; red crosses, outliers. \**p* < 0.05, \*\**p* < 0.005, \*\*\**p* < 0.0005; n.s., not significant.

**Supplementary Figure 6.**
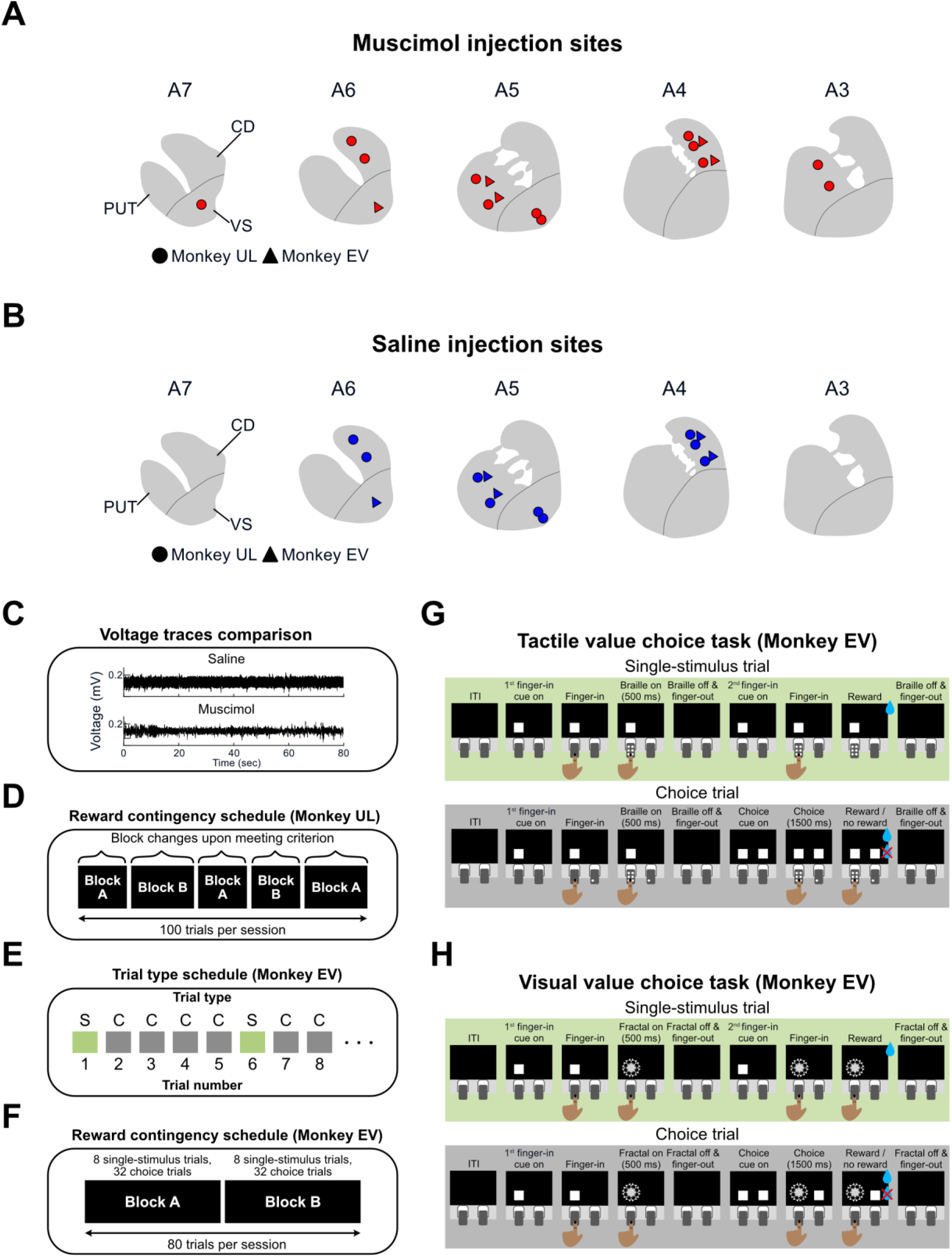
Injection sites, block schedules, and monkey-specific task variations in the inactivation study. **(A)** Muscimol injection sites for individual monkeys. **(B)** Saline injection sites for individual monkeys. **(C)** Voltage trace comparisons between muscimol and saline conditions. **(D)** Performance-based block schedule for Monkey UL. Block schedule in which reward contingencies were reversed according to performance criteria. Each box represents a single block. The criterion was dynamic, requiring 14 to 19 consecutive correct trials to trigger a reversal. **(E)** Trial sequence for Monkey EV. Schematic of the trial sequence, in which a single-stimulus trial was interleaved after every four choice trials throughout the session. S stands for single-stimulus trial and C stands for choice trial. **(F)** Fixed block schedule for Monkey EV. Block schedule in which reward contingencies were reversed after a fixed duration of 40 trials per block, independent of immediate performance. **(G-H)** Task variants for Monkey EV during inactivation study, consisting of single-stimulus trials and choice trials. **(G)** Tactile value choice task (T-VCT) variant for Monkey EV. **(H)** Visual value choice task (V-VCT) variant for Monkey EV.

**Supplementary Figure 7.**
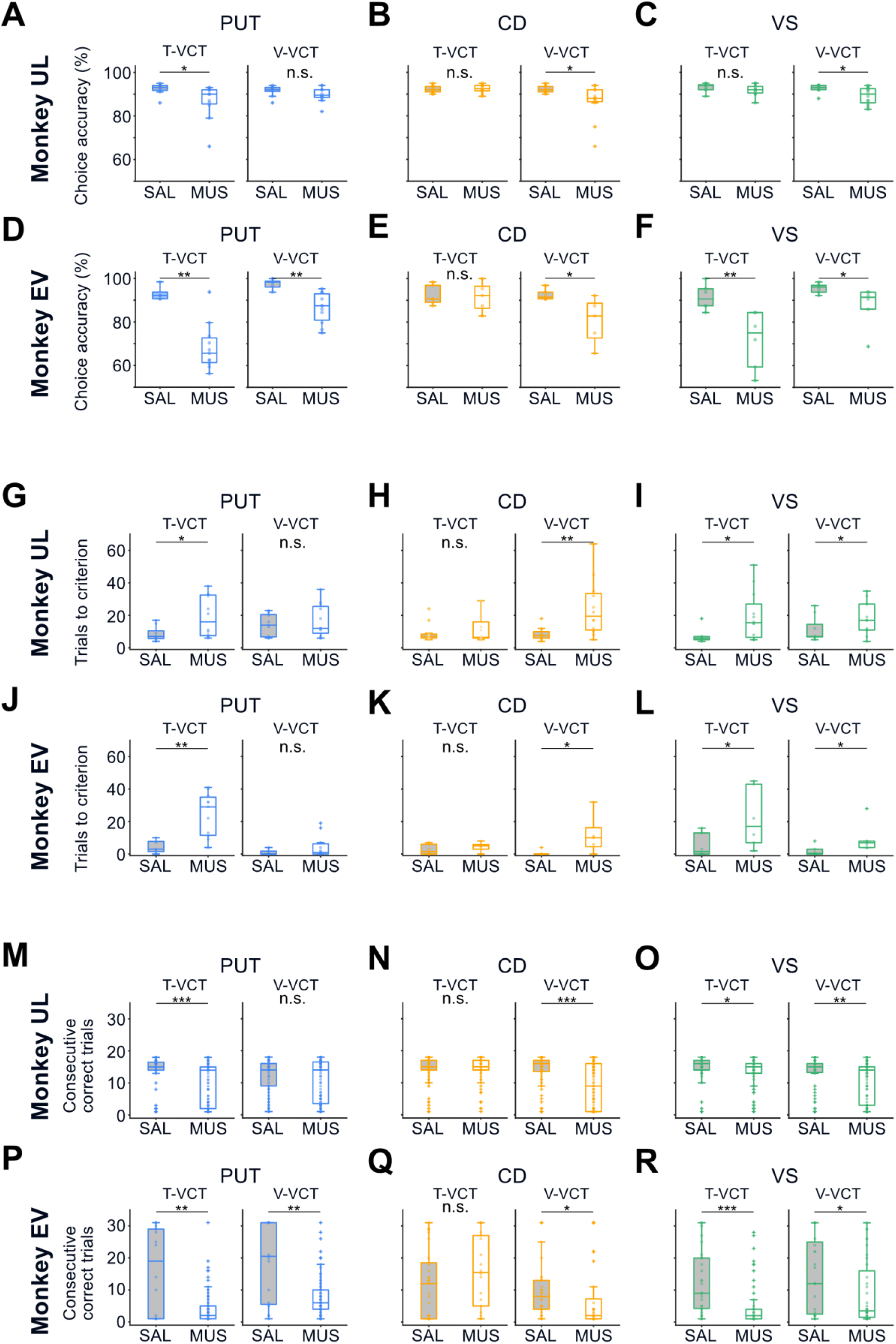
Individual behavioral performance during striatal inactivation. **(A-F)** Choice accuracy by subject. Comparison of correct performance rates under saline and muscimol conditions for each monkey. (A-C) Results for Monkey UL in the **(A)** putamen (T-VCT: one-tailed Wilcoxon rank-sum test, *p* = 0.0051; V-VCT: one-tailed Wilcoxon rank-sum test, *p* = 0.1012), **(B)** caudate (T-VCT: one-tailed unpaired t-test, *p* = 0.6758; V-VCT: one-tailed Wilcoxon rank-sum test, *p* = 0.0091), and **(C)** ventral striatum (T-VCT: one-tailed unpaired t-test, *p* = 0.1071; V-VCT: one-tailed Wilcoxon rank-sum test, *p* = 0.0126). (D-F) Results for Monkey EV in the **(D)** putamen (T-VCT: one-tailed Wilcoxon rank-sum test, *p* = 0.00275; V-VCT: one-tailed unpaired t-test, *p* = 0.002), **(E)** caudate (T-VCT: one-tailed unpaired t-test, *p* = 0.4282; V-VCT: one-tailed unpaired t-test, *p* = 0.0124), and **(F)** ventral striatum (T-VCT: one-tailed unpaired t-test, *p* = 0.00383; V-VCT: one-tailed Wilcoxon rank-sum test, *p* = 0.0108). **(G-L)** Trials to criterion by subject. Total trials required to reach the learning criterion. **(G-I)** Results for Monkey UL in the **(G)** putamen (T-VCT: one-tailed Wilcoxon rank-sum test, *p* = 0.0101; V-VCT: one-tailed unpaired t-test, *p* = 0.2641), **(H)** caudate (T-VCT: one-tailed Wilcoxon rank-sum test, *p* = 0.488; V-VCT: one-tailed Wilcoxon rank-sum test, *p* = 0.0021), and **(I)** ventral striatum (T-VCT: one-tailed Wilcoxon rank-sum test, *p* = 0.021; V-VCT: one-tailed Wilcoxon rank-sum test, *p* = 0.0367). **(J-L)** Results for Monkey EV in the **(J)** putamen (T-VCT: one-tailed unpaired t-test, *p* = 0.00286; V-VCT: one-tailed Wilcoxon rank-sum test, *p* = 0.1731), **(K)** caudate (T-VCT: one-tailed unpaired t-test, *p* = 0.1873; V-VCT: one-tailed Wilcoxon rank-sum test, *p* = 0.0152), and **(L)** ventral striatum (T-VCT: one-tailed Wilcoxon rank-sum test, *p* = 0.0455; V-VCT: one-tailed Wilcoxon rank-sum test, *p* = 0.0108). **(M-R)** Consecutive correct trials by subject. Total consecutive correct trials were counted without the first trial of each block. **(M-O)** Results for Monkey UL in the **(M)** putamen (T-VCT: one-tailed Wilcoxon rank-sum test, *p* < 0.0001; V-VCT: one-tailed Wilcoxon rank-sum test, *p* = 0.0921), **(N)** caudate (T-VCT: one-tailed Wilcoxon rank-sum test, *p* = 0.3826; V-VCT: one-tailed Wilcoxon rank-sum test, p < 0.0001), and **(O)** ventral striatum (T-VCT: one-tailed Wilcoxon rank-sum test, *p* = 0.015; V-VCT: one-tailed Wilcoxon rank-sum test, *p* = 0.001). **(P-R)** Results for Monkey EV in the **(P)** putamen (T-VCT: one-tailed Wilcoxon rank-sum test, *p* = 0.0049; V-VCT: one-tailed Wilcoxon rank-sum test, *p* = 0.00159), **(Q)** caudate (T-VCT: one-tailed Wilcoxon rank-sum test, *p* = 0.9269; V-VCT: one-tailed Wilcoxon rank-sum test, *p* = 0.0053), and **(R)** ventral striatum (T-VCT: one-tailed Wilcoxon rank-sum test, *p* < 0.0001; V-VCT: one-tailed Wilcoxon rank-sum test, *p* = 0.0274). Box plot elements: center line, median; box limits, 25th and 75th percentiles; whiskers, 1.5 x IQR; red crosses, outliers. *p < 0.05, **p < 0.005, ***p < 0.0005, n.s. not significant.

**Supplementary Figure 8.**
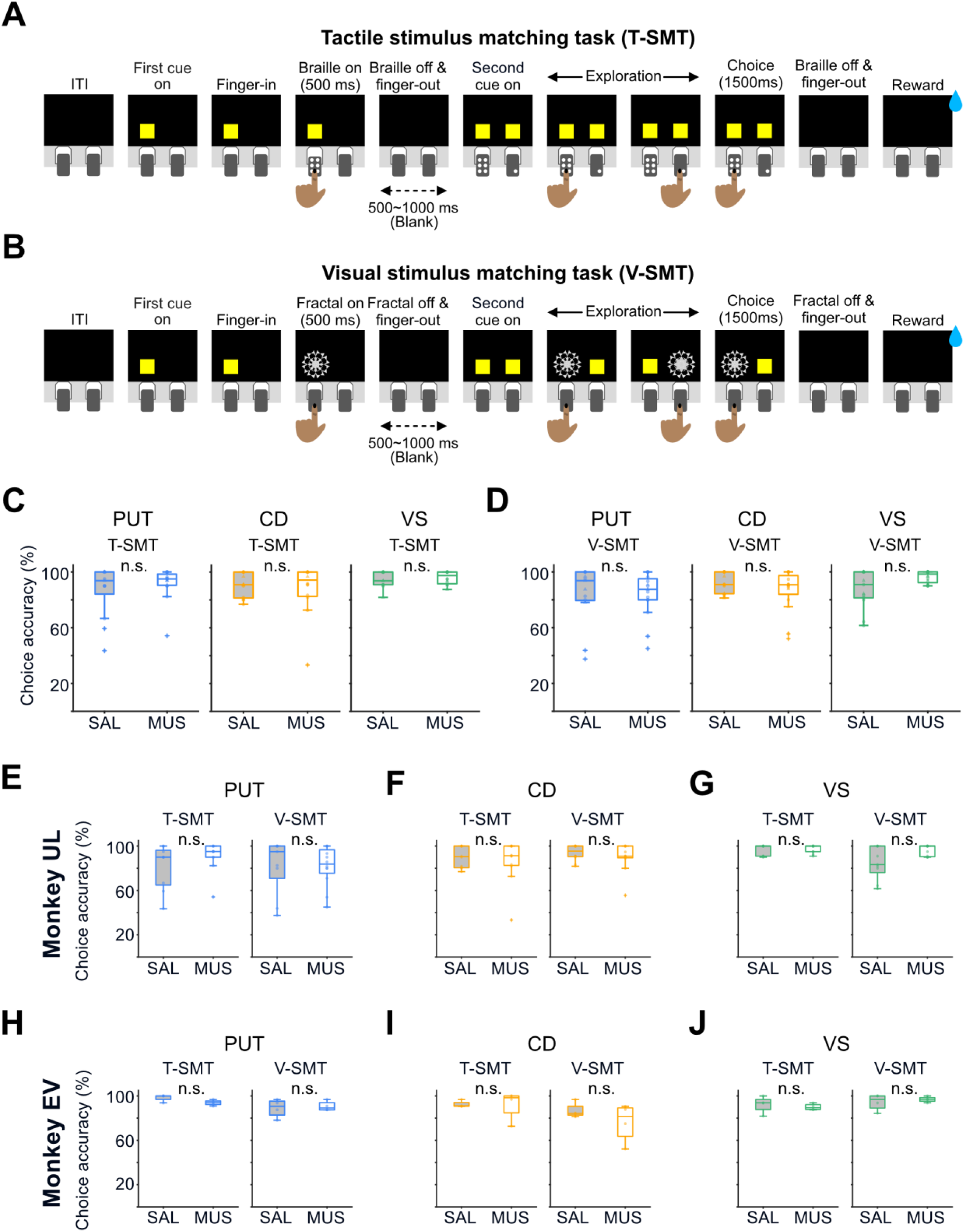
Behavioral performance in control matching tasks. **(A-B)** Task schematics. **(A)** Tactile Stimulus Matching Task (T-SMT). A delayed match-to-sample paradigm where monkeys were required to match a sample stimulus encountered at the beginning of the trial. **(B)** Visual Stimulus Matching Task (V-SMT). The procedure is identical to (A), except for the use of visual modality stimuli. **(C–D)** Correct performance rates in saline and muscimol conditions across the three striatal regions. **(C)** Performance comparison in T-SMT (Putamen: one-tailed Wilcoxon rank-sum test, *p* = 0.6724; caudate: one-tailed Wilcoxon rank-sum test, *p* = 0.6929; ventral striatum: one-tailed Wilcoxon rank-sum test, *p* = 0.6627). **(D)** Performance comparison in V-SMT (Putamen: one-tailed Wilcoxon rank-sum test, *p* = 0.3134; caudate: one-tailed Wilcoxon rank-sum test, *p* = 0.1938; ventral striatum: one-tailed Wilcoxon rank-sum test, *p* = 0.9534). **(E-G)** Matching task performance for Monkey UL. Comparison of correct performance in T-SMT (*left*) and V-SMT (*right*) under saline and muscimol conditions. **(E)** Putamen inactivation in T-SMT (one-tailed Wilcoxon rank-sum test, *p* = 0.91) and V-SMT (one-tailed Wilcoxon rank-sum test, *p* = 0.3188). **(F)** Caudate inactivation in T-SMT (one-tailed Wilcoxon rank-sum test, *p* = 0.6293) and V-SMT (one-tailed Wilcoxon rank-sum test, *p* = 0.2435). **(G)** Ventral striatum inactivation in T-SMT (one-tailed Wilcoxon rank-sum test, *p* = 0.8559) and V-SMT (one-tailed Wilcoxon rank-sum test, *p* = 0.9727). **(H-J)** Matching task performance for Monkey EV. Same conventions as in (E–G). **(H)** Putamen inactivation in T-SMT (one-tailed Wilcoxon rank-sum test, *p* = 0.06) and V-SMT (one-tailed unpaired t-test, *p* = 0.6303). **(I)** Caudate inactivation in T-SMT (one-tailed Wilcoxon rank-sum test, *p* = 0.857) and V-SMT (one-tailed unpaired t-test, *p* = 0.1583). **(J)** Ventral striatum inactivation in T-SMT (one-tailed unpaired t-test, *p* = 0.2954) and V-SMT (one-tailed unpaired t-test, *p* = 0.7148). Box plot elements: center line, median; box limits, 25th and 75th percentiles; whiskers, 1.5 x IQR; red crosses, outliers. \**p* < 0.05, \*\**p* < 0.005, \*\*\**p* < 0.0005; n.s., not significant.

**Supplementary Figure 9.**
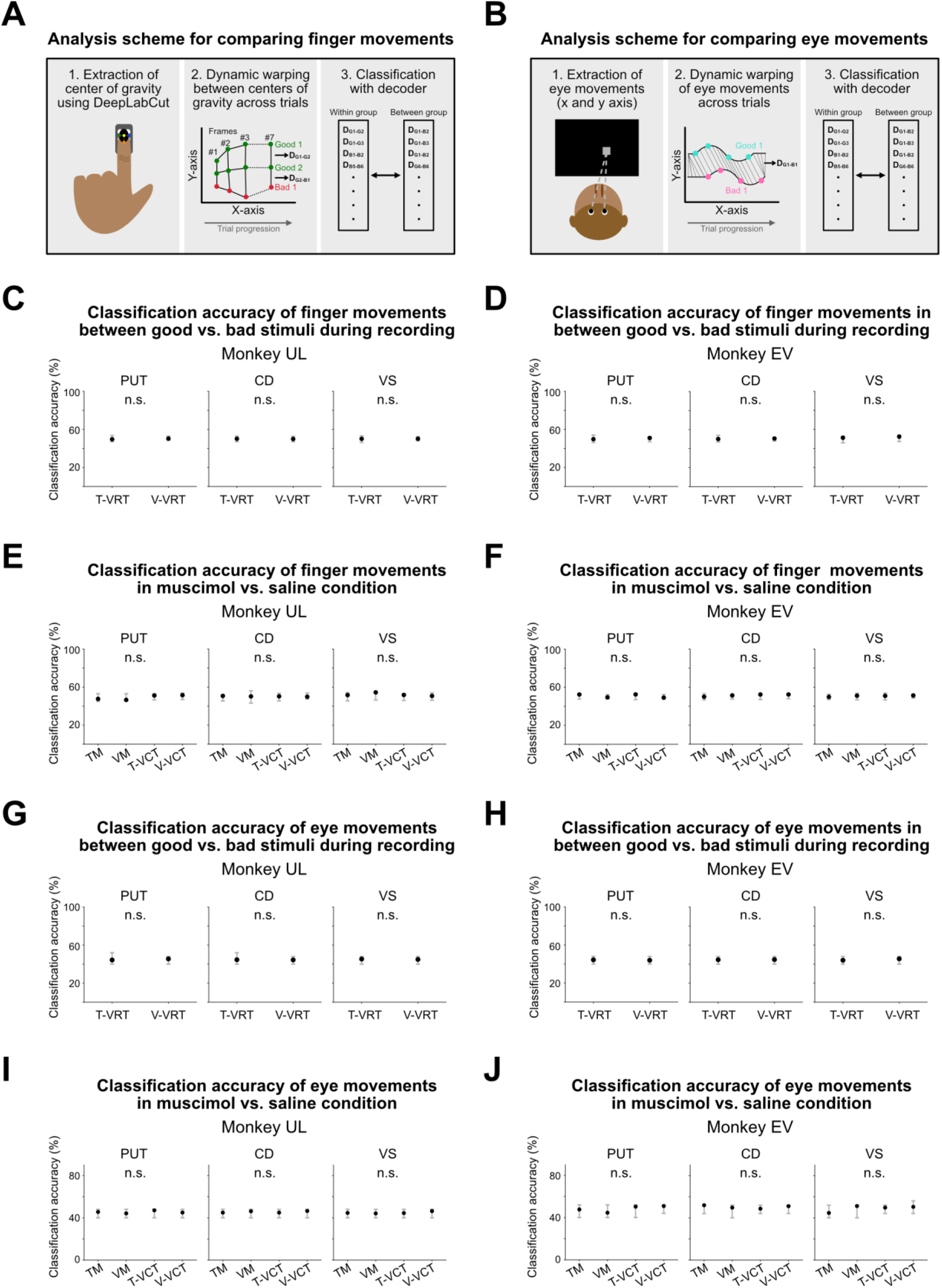
Analysis of finger and eye movements during recording and inactivation sessions. **(A)** Schematic illustration of the analysis of finger movements. The center of gravity for each tested frame was determined using three finger points extracted via DeepLabCut (left panel) as previously described (Hwang et al., 2025). Similarity of the center of gravity traces between trials was quantified using dynamic time warping (DTW) (middle panel). The classification decoder was trained and tested within the same condition and between different conditions (right panel). **(B)** Schematic illustration of the analysis of eye movements. Eye movements were extracted (left panel) and the similarity of eye traces between different trials was quantified using DTW (middle panel). The decoder was trained and tested using DTW distances within the same condition and between different conditions, and the corresponding performance was calculated (right panel). **(C-F)** The analysis of finger movements during recording sessions and inactivation sessions in two monkeys. **(C)** Classification accuracy of finger movements for Monkey UL during recording sessions in putamen, caudate, and ventral striatum. **(D)** Classification accuracy of finger movements for Monkey EV during recording sessions in putamen, caudate, and ventral striatum. The results indicate that the decoder failed to distinguish between distances within the same condition and those between different conditions based on tactile or visual value. The black indicators represent the mean classification accuracy, while the grayscale vertical bars with open circles indicate the 95% confidence interval of the null model (right-tailed z-test). **(E)** Classification accuracy of finger movements for Monkey UL during muscimol and saline sessions in putamen, caudate, and ventral striatum. **(F)** Classification accuracy of finger movements for Monkey EV during muscimol and saline sessions in putamen, caudate, and ventral striatum. **(G-J)** Analysis of eye movements during recording sessions and inactivation sessions in two monkeys. **(G)** Classification accuracy of eye movements for Monkey UL during recording sessions in putamen, caudate, and ventral striatum. **(H)** Classification accuracy of eye movements for Monkey EV during recording sessions in putamen, caudate, and ventral striatum. **(I)** Classification accuracy of eye movements for Monkey UL during muscimol and saline sessions in putamen, caudate, and ventral striatum. **(J)** Classification accuracy of eye movements for Monkey EV during muscimol and saline sessions in putamen, caudate, and ventral striatum. n.s., not significant.

**Table S1.** Classification of value-coding neurons.

| # | Stimulus period |  | Delay period |  | Period Type | Neuron Type | Analysis Method |
| --- | --- | --- | --- | --- | --- | --- | --- |
|  | Tactile | Visual | Tactile | Visual |  |  |  |
| 1 | X | X | X | X |  | Not a value neuron |  |
| 2 | X | X | X | O | Delay Period | Visual-selective value |  |
| 3 | X | X | O | X | Delay Period | Tactile-selective value |  |
| 4 | X | X | O | O | Delay Period | Bimodal value |  |
| 5 | X | O | X | X | Stimulus Period | Visual-selective value |  |
| 6 | X | O | X | O | *Based on ROC* | Visual-selective value | Use more extreme ROC |
| 7 | X | O | O | X | *Based on ROC* | Bimodal value | Use more extreme ROC |
| 8 | X | O | O | O | Delay Period | Bimodal value |  |
| 9 | O | X | X | X | Stimulus Period | Tactile-selective value |  |
| 10 | O | X | X | O | *Based on ROC* | Bimodal value | Use more extreme ROC |
| 11 | O | X | O | X | *Based on ROC* | Tactile-selective value | Use more extreme ROC |
| 12 | O | X | O | O | Delay Period | Bimodal value |  |
| 13 | O | O | X | X | Stimulus Period | Bimodal value |  |
| 14 | O | O | X | O | Stimulus Period | Bimodal value |  |
| 15 | O | O | O | X | Stimulus Period | Bimodal value |  |
| 16 | O | O | O | O | *Based on ROC* | Bimodal value | Use the most extreme ROC |

“O” indicates statistically significant value neural activity in the corresponding task and period
type (Wilcoxon rank-sum test). “X” indicates non-significant value neural activity in the
corresponding task and period type. “Based on ROC” indicates that neurons were classified
according to the more extreme ROC value across stimulus and delay periods, or the most
extreme ROC value between T-VRT and V-VRT. “Use more extreme ROC”: select the period
with the more extreme ROC within a modality-specific task. “Use the most extreme ROC”:
select the period with the most extreme ROC across T-VRT and V-VRT.

**Table S2.** Statistical details for the main figure.

| Figure | Detail | Statistics |
| --- | --- | --- |
| **1D** | T-VRT | Paired t-test, $p < 0.0001$ |
| | V-VRT | Paired t-test, $p < 0.0001$ |
| **3A-C** | Bimodal | Pearson's $\chi^2$ test, $\chi^2 = 5.1397$, $df = 2$, $p = 0.0765$ |
| | Tactile-selective | Pearson's $\chi^2$ test, $\chi^2 = 0.3804$, $df = 2$, $p = 0.8268$ |
| | Visual-selective | Pearson's $\chi^2$ test, $\chi^2 = 8.6857$, $df = 2$, $p = 0.013$; with Bonferroni post hoc tests: putamen vs. caudate, $p = 0.6537$; putamen vs. ventral striatum, $p = 0.1679$; caudate vs. ventral striatum, $p = 0.0096$ |
| **4C** | Bimodal, T-VRT | Kruskal-Wallis test ($H(2) = 2.59$, $p = 0.2743$). |
| | Bimodal, V-VRT | Kruskal-Wallis test ($H(2) = 1.11$, $p = 0.5728$). |
| **4D** | Tactile-selective | Kruskal-Wallis test ($H(2) = 0.000573$, $p = 0.9997$). |
| | Visual-selective | Kruskal-Wallis test ($H(2) = 0.0916$, $p = 0.9553$). |
| **4E** | Bimodal, T-VRT | Pearson's $\chi^2$ test, $\chi^2 = 3.5429$, $df = 2$, $p = 0.1701$ |
| | Bimodal, V-VRT | Pearson's $\chi^2$ test, $\chi^2 = 1.9859$, $df = 2$, $p = 0.3705$ |
| **4F** | Tactile-selective | Pearson's $\chi^2$ test, $\chi^2 = 6.3159$, $df = 2$, $p = 0.0425$ |
| | Visual-selective | Pearson's $\chi^2$ test, $\chi^2 = 0.1706$, $df = 2$, $p = 0.9182$ |
| **5B** | | Two-tailed Wilcoxon rank-sum test, $p < 0.0001$ |
| **5D-G** | Magenta outline | Permutation test; right-tailed z-test, $p < 0.05$ |
| **6E** | PUT, T-VCT | One-tailed Wilcoxon rank-sum test, $p < 0.0001$ |
| | PUT, V-VCT | One-tailed Wilcoxon rank-sum test, $p = 0.00152$ |
| **6F** | CD, T-VCT | One-tailed unpaired t-test, $p = 0.5183$ |
| | CD, V-VCT | One-tailed Wilcoxon rank-sum test, $p = 0.0007$ |
| **6G** | VS, T-VCT | One-tailed Wilcoxon rank-sum test, $p = 0.013$ |
| | VS, V-VCT | One-tailed Wilcoxon rank-sum test, $p = 0.00036$ |
| **6H** | PUT | Two-tailed Wilcoxon rank-sum test, $p < 0.0001$ |
| **6I** | CD | Two-tailed Wilcoxon rank-sum test, $p < 0.0001$ |
| **6J** | VS | Two-tailed Wilcoxon rank-sum test, $p = 0.5712$ |
| **7B** | PUT, T-VCT | One-tailed Wilcoxon rank-sum test, $p = 0.000154$ |
| | PUT, V-VCT | One-tailed Wilcoxon rank-sum test, $p = 0.4067$ |
| **7C** | CD, T-VCT | One-tailed Wilcoxon rank-sum test, $p = 0.3755$ |
| | CD, V-VCT | One-tailed unpaired t-test, $p = 0.000471$ |
| **7D** | VS, T-VCT | One-tailed Wilcoxon rank-sum test, $p = 0.00186$ |
| | VS, V-VCT | One-tailed Wilcoxon rank-sum test, $p = 0.00649$ |
| **7F** | PUT, T-VCT | One-tailed Wilcoxon rank-sum test, $p < 0.0001$ |
| | PUT, V-VCT | One-tailed Wilcoxon rank-sum test, $p < 0.0001$ |
| **7G** | CD, T-VCT | One-tailed Wilcoxon rank-sum test, $p = 0.8014$ |
| | CD, V-VCT | One-tailed Wilcoxon rank-sum test, $p < 0.0001$ |
| **7H** | VS, T-VCT | One-tailed Wilcoxon rank-sum test, $p < 0.0001$ |
| | VS, V-VCT | One-tailed Wilcoxon rank-sum test, $p < 0.0001$ |

**Table S3.** p-values for CCGP differences between adapted and unadapted trials.

| CCGP Differences |  | T-VRT |  | V-VRT |  |
| --- | --- | --- | --- | --- | --- |
| Subregions | Neuron type | Stim. | Delay | Stim. | Delay |
| PUT | Bimodal val. | 0.0336 | 0.0275 | 0.3523 | 0.2631 |
|  | Tactile-sel. val. | 0.0303 | 0.0078 |  |  |
|  | Visual-sel. val. |  |  | 0.0819 | 0.0463 |
| CD | Bimodal val. | 0.4566 | 0.2207 | 0.3701 | 0.1694 |
|  | Tactile-sel. val. | 0.8292 | 0.7377 |  |  |
|  | Visual-sel. val. |  |  | 0.0061 | 0.4796 |
| VS | Bimodal val. | 0.3974 | 0.3051 | 0.3727 | 0.2860 |
|  | Tactile-sel. val. | 0.0648 | 0.1499 |  |  |
|  | Visual-sel. val. |  |  | 0.4014 | 0.0974 |

The table presents p-values obtained by comparing the actual difference in cross-condition generalization performance (CCGP) between adapted and unadapted trials against a shuffled null distribution. Statistical significance was assessed using a right-tailed z-test. Dark gray shaded cells indicate statistical significance ( $p < 0.05$ ). Light gray shaded cells denote a strong trend toward significance ( $0.05 < p < 0.10$ ). Cells with diagonal lines indicate non-applicable comparisons, as modality-selective neurons were not analyzed during their non-preferred modality tasks. Abbreviations: PUT, putamen; CD, caudate; VS, ventral striatum; Stim., stimulus period.

**Table S4.**
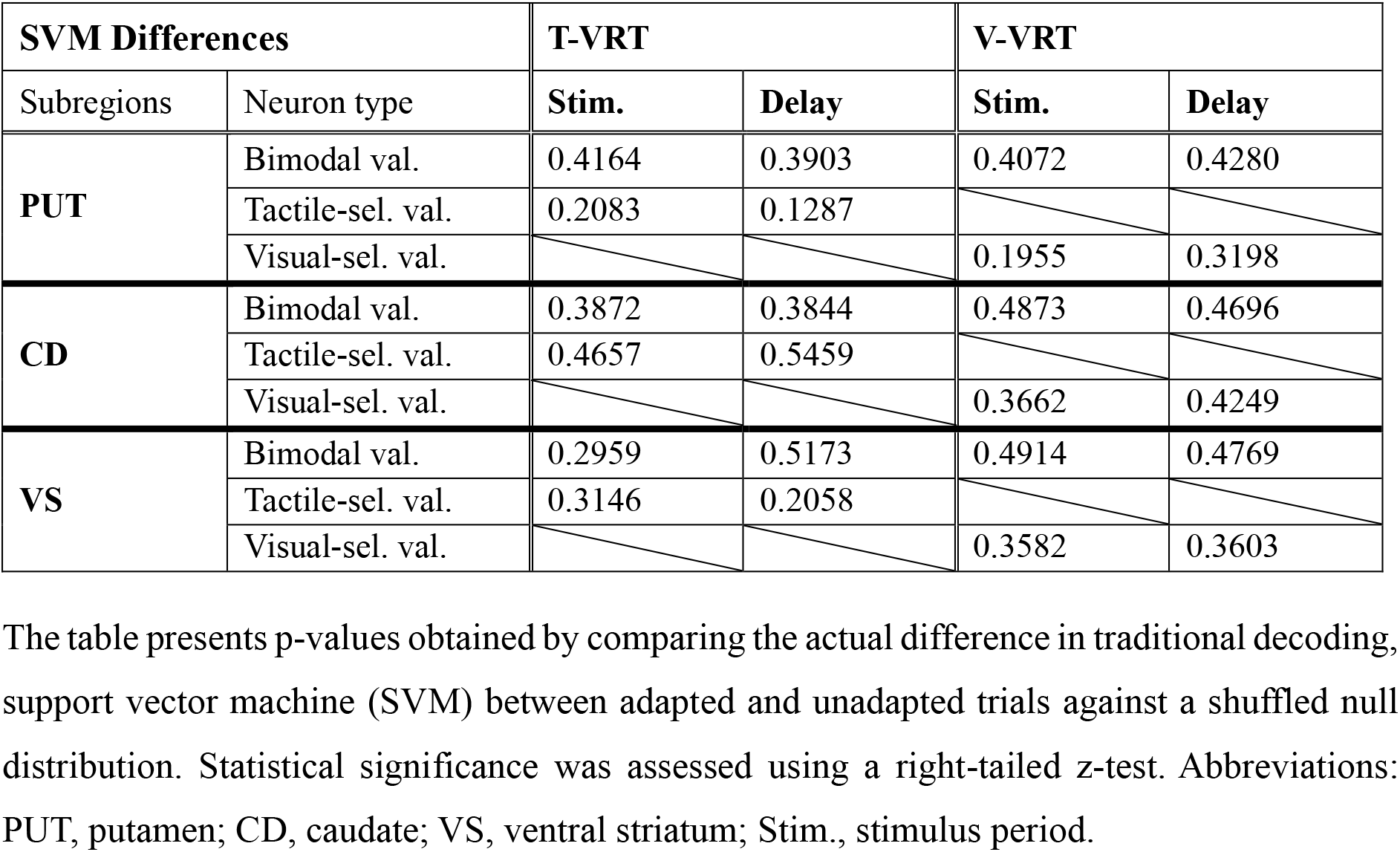
p-values for SVM differences between adapted and unadapted trials.

## Notes

### Competing Interest Statement

The authors have declared no competing interest.

